# Single cell mapping of the metabolic landscape of skin fibrosis in systemic sclerosis

**DOI:** 10.1101/2025.04.14.648761

**Authors:** Veda Devakumar, Yi-Nan Li, Tim Filla, Aleix Rius Rigau, Andrea-Hermina Györfi, Bilgesu Safak Tümerdem, Ranjana Neelagar, Minrui Liang, Christina Bergmann, Georg Schett, Jörg H.W. Distler, Alexandru-Emil Matei

## Abstract

Tissue resident cells undergo metabolic reprogramming during fibrotic tissue remodeling to meet their changing metabolic demands required for extracellular matrix production and phenotypic transitions in fibrosis. However, the metabolic reprogramming in fibrotic tissues has not yet been explored at single cell level with spatial resolution. Moreover, the spatial organization of metabolic niches in fibrotic tissues remains understudied. To address these gaps, we used imaging mass cytometry (IMC) and characterized the metabolic regulome, indicative of the activity in several key metabolic pathways in systemic sclerosis (SSc) as a prototypic systemic fibrotic disease. We identified a distinct metabolically active profile with high activity of glycolysis, TCA/OXPHOS, hypoxia and ROS signaling in fibroblasts, endothelial cells and macrophages in SSc patients with progressive skin fibrosis. These metabolic profiles are associated with expression of markers of profibrotic activation. Metabolically active fibroblasts might shape their microenvironment to induce a similar metabolic phenotype in neighboring endothelial cells and macrophages, facilitating profibrotic interactions. Consistently, specific interactions between metabolically defined, activated cell subsets are associated with the extent or progression of skin fibrosis. Thus, interfering with these metabolic niches might provide therapeutic opportunities in fibrotic diseases.

## Introduction

Systemic sclerosis (SSc) is a prototypical fibrotic disease characterized by excessive deposition of extracellular matrix (ECM) in the skin and internal organs, by autoimmunity and vasculopathy ^1^. Accumulating ECM often compromises organ function and results in high morbidity and mortality ^2^. Fibroblasts are the main effector cells that deposit excessive amounts of extracellular matrix to drive fibrotic tissue remodeling ^3–5^. Endothelial cells and macrophages are other cell types with key pathogenic roles in SSc ^2^. Endothelial cells can undergo endothelial-to-mesenchymal transition (EndMT) to contribute to the pool of collagen-producing myofibroblasts ^6–9^. Macrophages release profibrotic mediators to induce myofibroblast differentiation and collagen production to promote fibrotic tissue remodeling ^10,11^. Fibroblasts, endothelial cells and macrophages undergo metabolic reprogramming to meet the increased energy demands associated with ECM production, cellular proliferation and phenotypic transitions in fibrotic tissues ^12–14^. Fibroblasts and endothelial cells rely primarily on glycolysis and reduce fatty acid oxidation (FAO) in tissue fibrosis ^15–17^. Macrophages have divergent metabolic programs, with M1-polarized macrophages having a predominantly glycolytic phenotype with interruptions of tricarboxylic acid (TCA) cycle at several steps, while M2-polarized macrophages rely predominantly of FAO and OXPHOS and have an intact TCA cycle ^12^. However, these findings were obtained from isolated cells or from tissues without single cell resolution. Thus, previous studies did not characterize the heterogeneity of cellular metabolic profiles in tissue fibrosis, to understand the interplay of metabolically reprogrammed cells and identify local metabolic changes and metabolic niches.

Here, we aimed to characterize the distinct metabolic states of fibroblasts, endothelial cells and macrophages within their local niches in SSc skin, as well as associations of shifts in cellular metabolic phenotypes and their spatial relationships with clinical outcomes in SSc. We employed a recently described imaging mass cytometry (IMC)-based approach that allows simultaneous quantification of the expression of 22 key metabolic regulators (the “metabolic regulome”) in single cells with spatial resolution ^18^. These regulators included rate-limiting enzymes and metabolite transporters involved in main metabolic pathways such as glycolysis, TCA cycle, OXPHOS, FAO, fatty acid synthesis (FAS), pentose phosphate pathway (PPP) and intracellular amino acid transport, along with transcription factors involved in mitochondrial dynamics and with regulators of metabolic signaling pathways such as mTOR or AMPK pathway, hypoxia and ROS signaling. Analysis of the metabolic regulome can serve as a proxy for assessing the activity of the corresponding metabolic pathways ^18^.

## Materials and methods

### Study design

We performed IMC from skin biopsies of patients with SSc and normal donors, and analyzed 45340 single cells with spatial resolution, for which we simultaneously detected 39 markers. Our IMC antibody panel was designed to include antibodies against 12 phenotypic and activation markers (markers of epithelial cells, endothelial cells, T cells, macrophages, B cells and fibroblast activation markers such as αSMA and FAP), 22 metabolic markers (to characterize cellular metabolic states), and 5 structural markers (for tissue segmentation) (Table S1).

### Study participants

All SSc patients (n = 9) fulfilled the American College of Rheumatology (ACR) / European Alliance of Associations for Rheumatology (EULAR) 2013 criteria. Normal donors (n = 7) were matched for age and sex served as controls. Skin biopsies of SSc patients were taken at the forearm 15 ± 2 cm proximal from the styloid process of the ulna using disposable biopsy punches (pfm medical, Cologne, Germany / Gifu, Japan). Demographical and clinical features of the patients with SSc are presented in Table S2.

Progressive skin fibrosis was defined as ≥ 25% increase of modified Rodnan skin score (mRSS) in the past 12 months. Early SSc was defined as less than 3 years from the onset of the first non-Raynaud’s symptom. Skin biopsies from controls were collected from patients undergoing surgery either for trauma (without any injury at the region where the biopsy was taken from), for degenerative joint disease (osteoarthritis), or for cosmetic surgery.

The study was performed in accordance with the Declaration of Helsinki. Ethical approval was obtained by the ethical committees of the Medical Faculties of the Universities of Düsseldorf and Erlangen. All patients and controls signed an informed consent form approved by the respective ethical committees.

### Antibody validation, conjugation and titration

Antibody validation was performed as previously described ^9,19,20^. Antibodies that were not available as conjugates with rare metal isotopes were purchased as carrier-free purified antibodies. These antibodies were initially evaluated by standard immunofluorescence (IF) staining on FFPE sections of skin and titrated in IF experiments to identify optimal dilutions. Only antibodies that demonstrated the expected staining pattern (based on literature) in IF and that had a good signal-to-background ratio (on visual evaluation) were used for subsequent steps. These antibodies were conjugated with lanthanide (Ln) metal isotopes using the Maxpar X8 metal conjugation kit (Standard Biotools, #201300) following manufacturer’s protocol. Afterwards, metal-labeled antibodies were diluted to stock concentrations of 0.5 mg/ml in Antibody Stabilizer PBS (Boca Scientific, #131050) supplemented with 0.05% sodium azide.

In a second validation step, all metal-labeled antibodies were tested in IMC. Serial antibody dilutions were tested in consecutive FFPE sections of human skin. The antibodies were mixed with phosphate buffered saline (PBS) with 0.5% bovine serum albumin BSA at their respective optimal dilutions to generate an antibody cocktail for subsequent immunolabelling of all tissue sections.

### Tissue preparation and staining

Five-micron thick formalin-fixed paraffin-embedded (FFPE) sections were stained with the antibody cocktail containing all 39 antibodies. The slides were incubated at 65[for 1 hour in a dry oven, followed by deparaffinization in fresh xylene and rehydration through a graded ethanol series from 100% to 70%. The sections were then transferred to double-distilled water (ddH_2_O) followed by PBS. Antigen retrieval was conducted in preheated Tris Buffer (10 mM Tris, 0.05% Tween 20, pH 10.0) at 96°C for 30 minutes using an Eppendorf ThermoMixer with a 50 ml block, without shaking, followed by a 20-minute cooldown at room temperature. Non-specific binding was blocked with 2% BSA in PBS for 1 hour at room temperature.

For each section, the antibody cocktail, prepared in a total volume of 100 µL in PBS with 0.5% BSA, was applied, and the tissue sections were incubated with the antibody cocktail overnight at 4[in a humidified set up. Following a washing step with PBS with 0.1% Tween 20 Detergent (PBST) and two additional washing steps with PBS, the sections were stained with a nuclear intercalator solution containing Iridium 193 and 191 isotopes (dilution 1:400, Standard Biotools, #201192A) for 5 min at room temperature. After another washing step, samples were then air-dried for subsequent IMC acquisition. Hematoxylin and eosin (H&E) were prepared from a consecutive section and imaged using a Nanozoomer S60 slide scanner.

### IMC data acquisition

Data were acquired on a Helios time-of-flight mass cytometer coupled to a Hyperion Imaging System (Standard Biotools). Optical images of the slides were obtained before laser ablation using the Hyperion software. Regions of interest (ROI) included the three major layers of the skin, epidermis, dermis and hypodermis at comparable ratios to each other. The machine underwent calibration, and a daily quality control check was conducted using a tuning slide spiked with three metal elements (Standard Biotools, #201088) following the manufacturer’s instructions. Laser ablation was carried out at a frequency of 200 Hz, resulting in a pixel size of 1 µm^2^. All IMC data were saved in .mcd and .txt file formats. The Standard Biotools CyTOF software (Version 7.0.8493, Standard Biotools) was used for acquisition and data export.

### Workflow of IMC data processing, visualization and analysis

For quality control, individual .mcd files were reviewed independently by two scientists, confirming the staining quality in each channel across all acquired images. For visualization, single- or multiple-channel 16-bit .tiff files were exported from MCD Viewer (Version 1.0.560.6, Standard Biotools), and pseudocolored images were prepared from .tiff images using ImageJ / Fiji software (Version 1.53c, https://fiji.sc). Furthermore, for each antibody staining, the relationship between the signal intensity with the signal-to-noise ratio was computed and visualized.

Initial data pre-processing, including segmentation and generation of single-cell data was performed using the *Steinbock* pipeline (https://github.com/BodenmillerGroup/steinbock ^21^). Cell segmentation was carried out with the *DeepCell/Mesmer* deep learning algorithm, with Histone and Ir as nuclear markers, and PDGFRA, CD31, CD45, CD20, CD3, CD68, as well as ICSK1and ICSK2 from the Segmentation kit as cytoplasmatic markers ^22^.

The single-cell data generated with the Steinbock pipeline included mean signal intensity within each cell and spatial coordinates for cellular localization. Poorly segmented non-cellular objects were removed from the dataset using the thresholds of area < 20 or > 250 pixels. The *imcRtools* R package was used to load the data and construct a *SpatialExperiment* (spe) R object ^21,23^. An inverse hyperbolic sine (arcsinh) transformation with a coefficient of 1 was applied to all the datasets. The arcsinh transformed data were normalized by computing a z-score with the entire dataset as reference.

Subsequently, gating was performed to identify endothelial cells (positive for CD31), macrophages (positive for CD45 and CD68), other immune cells (positive for CD45 and negative for CD68), epithelial cells (positive for E-Cadherin) and fibroblasts (negative for CD31, CD45 and E-Cadherin). All the subsets identified were positive for Histone3. The gating process and the inspection of spatial distribution of the gated cells was carried out with the *cytomapper* R package ^24^.

### Glycolysis score and tricarboxylic acid / oxidative phosphorylation score computation

A glycolysis score was computed as the average expression of the key glycolytic enzymes GLUT1, HK1, PFKL1, PKM2, LDHA and GAPDH, as described ^18^. Similarly, a tricarboxylic acid (TCA) / oxidative phosphorylation (OXPHOS) score was computed as the average expression of CS, OGDH, SDHA and ATP5A ^18^.

### Single-cell clustering and dimensionality reduction

The populations identified by gating were analyzed individually using dimensionality reduction (uniform manifold approximation and projection, UMAP) and clustering (Rphenograph) with the *imcRtools* R package. The z-rescaled expressions of the metabolic markers, i.e. GLUT1, HK1, PFKL1, PKM2, LDHA, GAPDH, CS, OGDH, SDHA, ATP5A, CD98, ACAC, CPT1a, G6PD, p-mTOR, p-AMPK, NOX4, HIF1α, PGC-1α, TFAM and p-NRF2 were employed for dimensionality reduction and clustering. Clusters at different k neighbor values were evaluated using UMAP and protein expression patterns, and final cluster definition was selected using a k value that yielded clusters with high cluster stability (high average silhouette width and neighborhood purity) that preserved the highest number of biologically meaningful metabolic profiles (k = 500 for fibroblasts and endothelial cells, k = 100 for macrophages). These cellular subsets were annotated according to the expression levels of specific markers and pathways.

### Neighborhood analysis

Spatial analysis was carried out using the *imcRtools* R package ^21^. Adjacent cells were first identified by Delaunay triangulation, with distance thresholds of 80 µm between their centroids. Subsequently, cellular neighborhoods (CNs) were identified as previously described ^9,20^. In brief, the composition of neighboring cells was calculated for each cell, and CNs were identified by K-means clustering of these compositions. Thus, the cells within one CN were similar with regards to the types and frequencies of their neighboring cells. Visualization of cell composition within CNs was performed by heatmaps rescaled to illustrate the frequency of each cell type within the respective CN relative to its frequency in the entire dataset, unless otherwise specified.

### Pairwise interaction analysis

To analyze what cells are found in spatial proximity more frequently than expected by chance, we performed cellular pairwise interaction analysis using the *imcRtools* R package ^21^. Permutation testing was employed to compare the neighboring cells of each cell type for each ROI individually to an empirical null distribution obtained by randomly shuffling cell type labels for 1000 permutations. A score was then calculated by averaging the test results from each ROI, as described ^20,25,26^. Dot plots and interaction network visualizations of the average scores for all ROIs were generated using *ggplot2* and *ggraph* R packages.

### Profiling of distances between different cell subsets

Quantification of the distances between the centroids of specific cells was carried out using the *imcRtools* R package. The minimum distances of different cell subsets to reference cells were plotted as density plots using density estimates using Gaussian kernels. Different cell subsets were annotated as belonging to niches of reference cell types if they were present within 40 µm^2^ from the reference cells.

### Data visualization and statistics

Quantitative data are shown either as bar graphs, violin plots, scatter plots or heatmaps. Data in bar graphs are presented as median ± interquartile range (IQR) with individual data points plotted as dots. Violin plots visualize the distribution of the data and its probability density, along with individual numbers labeled, as well as the median and quartiles of the data. Statistical significance was calculated with custom R scripts. Unless stated otherwise, Mann-Whitney U non-parametric testing was used for all two-group comparisons and Kruskall-Wallis with Dunn’s post-hoc test was used for the comparisons involving more than two groups. Relationships between continuous variables were assessed using non-parametric Spearman correlation. p-values < 0.05 were considered statistically significant.

## Results

### Progressive skin fibrosis in SSc is characterized by a distinct metabolic regulome of dermal cells

We characterized the metabolic regulome of the skin of SSc patients at single-cell level using an IMC antibody panel with 22 metabolic markers along with 19 markers to identify cellular subsets and structural markers for tissue segmentation (Fig. 1, Fig. S1, Table S1). We identified a total of 45,340 single cells by IMC (23,643 cells from SSc patients and 21,697 cells from controls).

**Figure 1:**
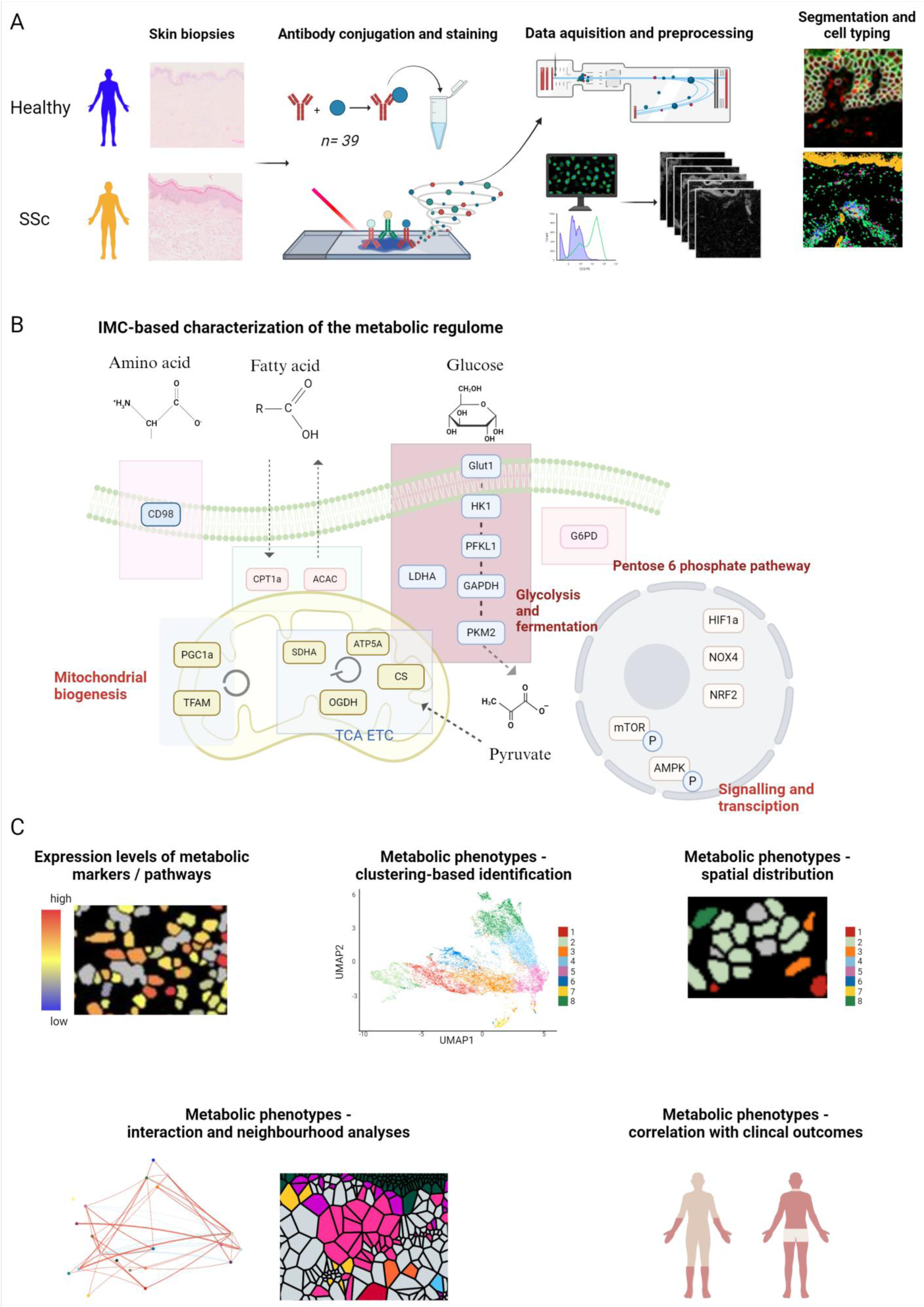
Schematic representation of the IMC workflow. **A.** Schematic representation of the IMC workflow starting from biopsies from SSc donors and controls. The antibodies were conjugated to heavy metal isotopes to stain the biopsies for their use in IMC. Upon laser ablation, time of flight of the metals were calculated to provide the expression level of each marker for each 1 µm^2^ pixel. Single cells were identified by segmentation, and identification of main cell populations (fibroblasts, endothelial cells, macrophages and keratinocytes) by gating according to expression of E-Cadherin, CD45, CD68 and CD31 was subsequently performed. **B.** Schematic representation of the markers included for evaluation of the metabolic regulome and the pathways they are involved in. **C.** Schematic representation of the analyses performed after gating: analysis of expression level of metabolic markers, identification of metabolically defined cell subsets by clustering using the metabolic markers, with subsequent evaluation of the spatial relationships, cell-cell interaction and cellular neighborhoods. Metabolic phenotypes were then correlated with clinical outcomes. This figure was created with Biorender.com. IMC: imaging mass cytometry; SSc: systemic sclerosis; UMAP: Uniform Manifold Approximation and Projection.

We first evaluated the expression profile of the metabolic regulators in dermal cells (defined as E-Cadherin^-^) in SSc and control skin. We observed that SSc patients with progression of skin fibrosis at the time of biopsy had a distinct metabolic regulome that clearly differed from the metabolic regulome of normal donors. This metabolic regulome in SSc patients with progressive skin fibrosis was characterized by the highest levels of expression of key metabolic regulators for glycolysis, TCA/OXPHOS, PPP, hypoxia and ROS signaling (Fig. 2A, B). In contrast, SSc patients with stable skin fibrosis had a metabolic regulome relatively similar to controls (Fig. 2A, B).

**Figure 2:**
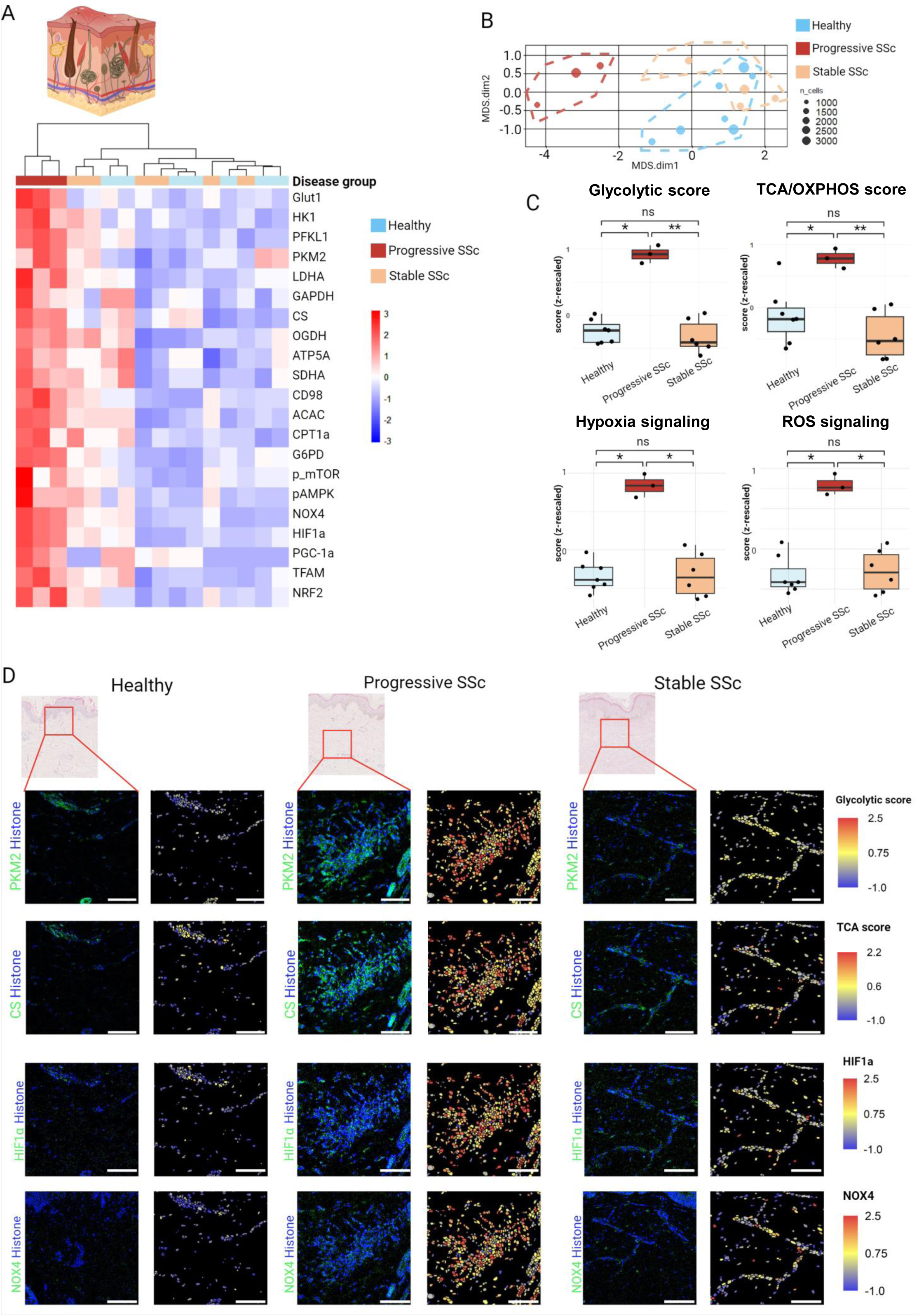
Distinct metabolic regulome of dermal cells in progressive skin fibrosis in SSc. **A.** Expression of metabolic markers across the groups: progressive SSc, stable SSc or controls, shown as heatmap, and illustration highlighting that the analyses were made on all dermal cells. **B.** MDS plot illustrating a distinct expression profile of the metabolic markers in dermal cells of SSc patients with progressive skin fibrosis than patients with stable skin fibrosis or controls. **C.** Glycolysis score (calculated by averaging the expression of enzymes belonging to the glycolysis pathway: GLUT1, HK1, PFKL1, PKM2, LDHA, GAPDH), TCA/OXPHOS score (calculated by averaging the expression of enzymes belonging to the TCA cycle and OXPHOS: CS, OGDH, ATP5A, SDHA), hypoxia (expression of HIF1α) and ROS (expression of NOX4) signaling in dermal cells across the groups: progressive SSc, stable SSc and controls, shown as box plot. **D.** Representative images illustrating spatial distribution of the glycolytic and TCA/ETC scores, and of the hypoxia and ROS signaling plotted on cell segmentation masks along with corresponding HE images and images showing expression of PKM2 (a glycolytic enzyme), CS (an enzyme from the TCA cycle), HIF1α (hypoxia signaling) and NOX4 (ROS signaling) at pixel level in dermal cells across the groups: progressive SSc, stable SSc and controls. Scale bars: 25 µm. Statistical significance was determined by the Kruskal Wallis test with Dunn’s post-hoc test. 0.05>*P*>0.01*, 0.01>*P*>0.001**, *P*<0.001***. This figure was created with Biorender.com. SSc: systemic sclerosis; TCA: tricarboxylic acid; OXPHOS: oxidative phosphorylation; GLUT1: Glucose transporter 1; HK1: Hexokinase1; PFKL1: Phosphofructokinase Ligand 1; PKM2: Pyruvate kinase isoform M2; LDHA: Lactate dehydrogenase A; GAPDH: Glyceraldehyde 3-phosphate dehydrogenase; CS: Citrate synthase; OGDH: 2-oxoglutarate dehydrogenase; ATP5A: Mitochondrial adenosine triphosphatase alpha chain; SDHA: succinate dehydrogenase complex; NOX4: Nicotinamide adenine dinucleotide phosphate hydrogen oxidase 4; ROS: Reactive oxygen species; HE: hematoxylin and eosin.

We further computed a glycolytic score and a TCA/OXPHOS score, which reflect the activity of the respective pathways ^18^. The glycolytic and the TCA/OXPHOS scores were highly increased in SSc patients with progressive skin involvement compared to those with stable/regressive skin fibrosis and to controls (Fig. 2C, D). The glycolytic score did not show any associations with other clinical features, whereas the TCA/OXPHOS score was numerically lower in patients with late disease (Fig. S2). Furthermore, SSc patients with progressive skin involvement had higher levels of hypoxia and ROS signaling than normal skin samples or skin from SSc patients with stable skin involvement (Fig. 2C, D). SSc patients with progressive skin fibrosis thus show a distinct metabolic phenotype independent of other manifestations that distinguishes them from non-progressors.

### Distinct metabolic profiles of fibroblasts subsets

We next aimed to characterize the metabolic regulome of the individual cell subpopulations in human skin (identified by gating, see Fig. S3) based on single cell analyses. We performed unbiased clustering of fibroblasts according to the metabolic markers. We identified eight distinct metabolic phenotypes that we termed “metabolically defined fibroblast (Fib-MET) phenotypes” (Fig. 3A, B). We annotated these metabolically defined phenotypes according to their expression levels of respective metabolic markers and pathways: 1) TCA/OXPHOS^hi^PGC-1α^hi^_Fib, expressing high levels of the TCA/OXPHOS enzymes and of PGC-1α, and lower levels of glycolytic enzymes, suggesting alternate metabolite sources other than glucose for TCA; 2) Met^hi^_Fib, expressing the highest levels of glycolytic enzymes and of TCA/OXPHOS enzymes, of the key FAO enzyme CPT1A and of hypoxia (HIF1α) and ROS synthesis capacity (NOX4), demonstrating a highly metabolically active phenotype; 3) TCA/OXPHOS^int^PGC-1α^hi^_Fib, with a similar expression pattern as TCA/OXPHOS^hi^PGC-1α^hi^_Fib, but with a markedly lower expression levels of the TCA/OXPHOS enzymes; 4) pmTOR^hi^CD98^hi^_Fib, with a high activity of the mTOR pathway and upregulation of the amino acid transporter CD98; 5) Met^low^_Fib, expressing the lowest levels of most metabolic regulators, including levels of enzymes involved in glycolysis and TCA/OXPHOS, thus suggesting a metabolically inactive, resting phenotype; 6) Glycolysis^int^CS^hi^OGDH^low^_Fib, that express high levels of CS, but low levels of OGDH, suggesting usage of TCA metabolites in cataplerotic reactions; 7) Glut1^hi^_Fib, expressing high levels of Glut1, but low levels of other glycolysis enzymes, indicating high glucose uptake, but usage in other processes than glycolysis, and 8) Glycolysis^int^PGC-1α^low^FAO^low^_Fib, indicating usage of glucose in glycolysis and PPP pathways with low levels of OXPHOS and FAO, and reduced mitochondrial biogenesis (Fig. 3B-F).

**Figure 3:**
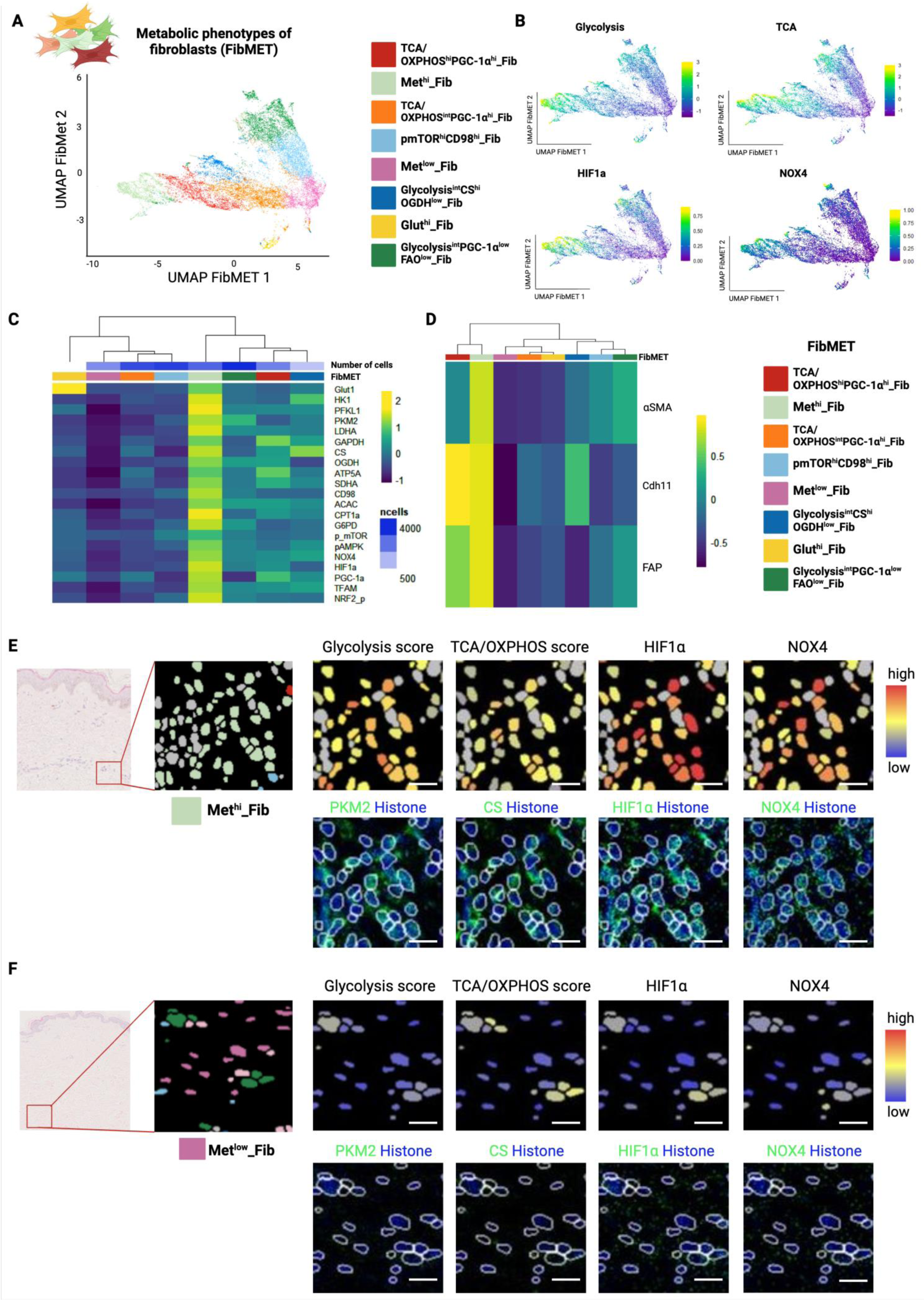
Identification of metabolically defined populations of fibroblasts. **A.** UMAP plots generated based on expression of metabolic markers and colour coded by metabolically defined fibroblast subsets, and illustration of the distinct fibroblast subsets. **B.** Glycolysis score (calculated by averaging the expression of enzymes belonging to the glycolysis pathway: GLUT1, HK1, PFKL1, PKM2, LDHA, GAPDH), TCA/OXPHOS score (calculated by averaging the expression of enzymes belonging to the TCA cycle and OXPHOS: CS, OGDH, ATP5A, SDHA), hypoxia (expression of HIF1α) and ROS (expression of NOX4) signaling in fibroblasts plotted on UMAP. **C.** Expression of metabolic markers across metabolically defined fibroblast subsets, plotted as heatmap. **D.** Expression of αSMA, Cadherin 11 and FAP across metabolically defined fibroblast subsets, plotted as heatmap. **E-F.** Representative images illustrating spatial distribution of the Met^hi^_Fib (E) and of Met^low^_Fib (F), along glycolysis and TCA/ETC scores and hypoxia and ROS signaling, plotted on cell segmentation masks, and images showing expression of PKM2 (a glycolytic enzyme), CS (an enzyme from the TCA cycle), HIF1α (hypoxia signaling) and NOX4 (ROS signaling) at pixel level. Corresponding HE images were included. Scale bars: 25 µm. This figure was created with Biorender.com. UMAP: Uniform Manifold Approximation and Projection; HE; hematoxylin and eosin; Fib: fibroblasts; TCA: tricarboxylic acid; OXPHOS: oxidative phosphorylation; GLUT1: Glucose transporter 1; HK1: Hexokinase1; PFKL1: Phosphofructokinase Ligand 1; PKM2: Pyruvate kinase isoform M2; LDHA: Lactate dehydrogenase A; GAPDH: Glyceraldehyde 3-phosphate dehydrogenase; CS: Citrate synthase; OGDH: 2-oxoglutarate dehydrogenase; ATP5A: Mitochondrial adenosine triphosphatase alpha chain; SDHA: succinate dehydrogenase complex; NOX4: Nicotinamide adenine dinucleotide phosphate hydrogen oxidase 4; ROS: Reactive oxygen species; FAP: fibroblast activation protein.

### Metabolically defined fibroblast populations differ in the expression levels of activation markers

We observed a distinct expression profile of markers of fibroblasts subsets and/or fibroblast activation markers αSMA, FAP and CDH11 among the Fib-MET subsets, indicating that fibroblast subsets defined by their metabolic profiles can differ with regards to their expression of phenotypic markers (Fig. 3D). The Met^hi^_Fib expressed the highest levels of αSMA and FAP and displayed thus a myofibroblast phenotype with a high pro-fibrotic potential.

On the other hand, several Fib-MET subsets such as TCA/OXPHOS^hi^PGC-1α^hi^_Fib and Glycolysis^int^PGC-1α^low^FAO^low^_Fib showed comparable levels of phenotypic markers such as αSMA and FAP and could thus not be distinguished by phenotypic markers alone (Fig. 3D). This suggests that the metabolic phenotyping of fibroblasts can reveal additional subpopulations of fibroblasts beyond those defined by classical markers.

### Changes in frequencies of metabolically distinct fibroblast subsets in SSc patients with progressive skin involvement

We next evaluated the changes in frequency of the Fib-MET phenotypes in the skin of SSc patients. Since the dermal cells of SSc patients with progressive skin fibrosis have a distinct metabolic regulome than SSc patients with stable skin fibrosis and controls, we hypothesized that the dermal fibroblasts of SSc patients with progressive skin fibrosis might demonstrate more pronounced changes in the frequencies of Fib-MET subsets. Indeed, we observed profound changes in the frequencies of many fibroblast subsets SSc patients with progressive skin fibrosis when compared to those with stable skin fibrosis and with controls (Fig. 4 and Fig. S4). Of these, the most prominent changes were the increase in the frequency of the Met^hi^_Fib and the decrease in the frequency of Met^low^_Fib in SSc patients with progressive skin fibrosis compared to patients with stable skin fibrosis and with controls (Fig. 4B-E). The fibroblasts in SSc patients with progressive skin fibrosis are metabolically polarized towards a Met^hi^_Fib phenotype, with more than 70% of all fibroblasts belonging to this phenotype, and almost no fibroblasts belonging to the Met^low^_Fib phenotype (Fig. 4). The TCA/OXPHOS^hi^PGC-1α^hi^_Fib were numerically increased in SSc patients with progressive skin fibrosis, whereas other Fib-MET populations such as pmTOR^hi^CD98^hi^_Fib, Glut^hi^_Fib or Glycolysis^int^PGC-1α^low^FAO^low^_Fib were numerically increased in patients with stable or regressive skin fibrosis compared to patients with progressive skin fibrosis or controls (Fig. 4B). These findings provide evidence for a profound metabolic reprogramming of fibroblasts in SSc patients with progression of skin fibrosis, with a possible differentiation of the Met^low^_Fib into Met^hi^_Fib.

**Figure 4:**
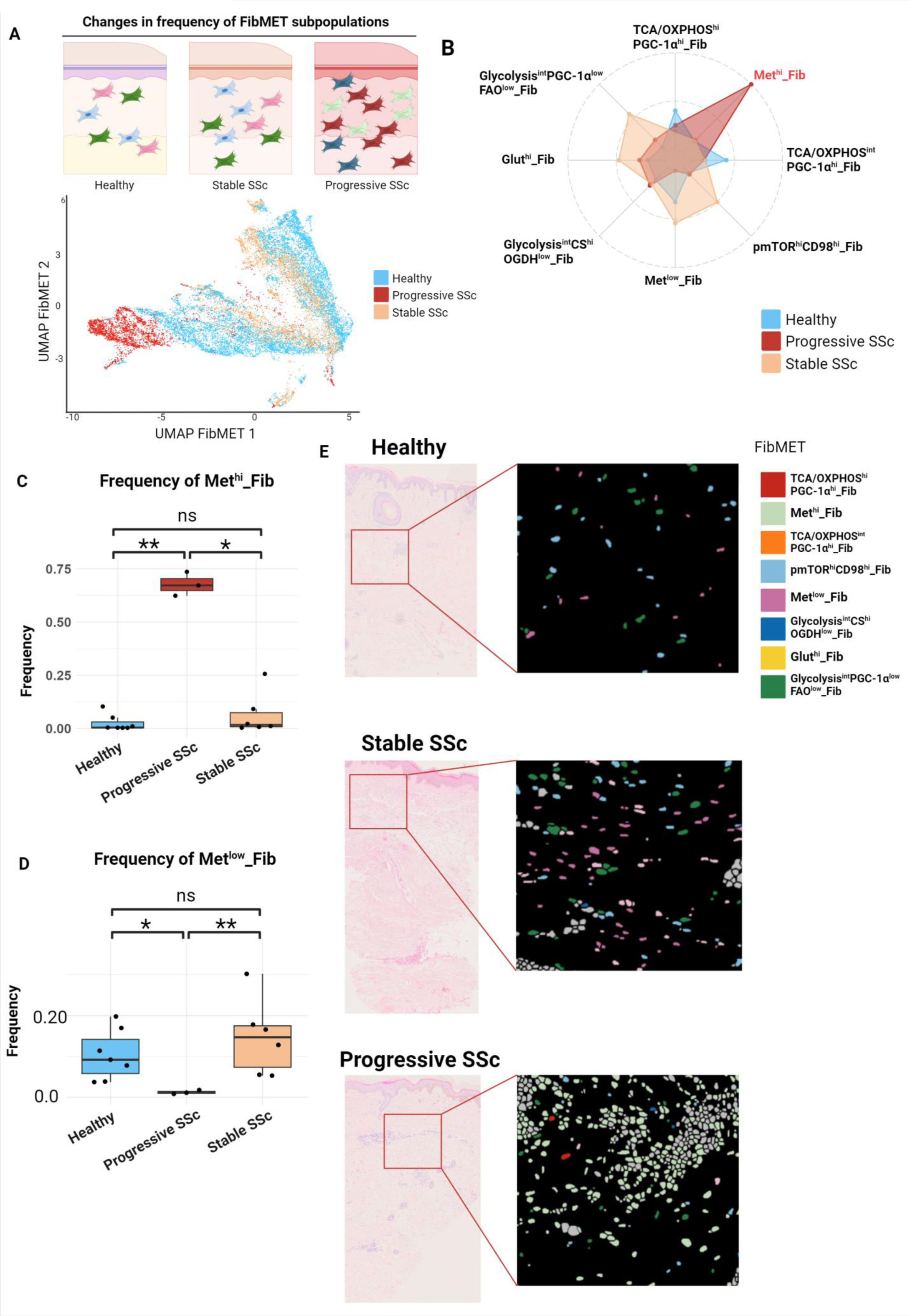
Shifts in frequencies of metabolically defined fibroblast subpopulations in SSc patients with progressive skin fibrosis. **A.** UMAP plots generated based on expression of metabolic markers and colour coded by the groups: progressive SSc, stable SSc or controls, and illustration of the shifts in fibroblast subsets across these groups. **B.** Radar chart illustrating the shifts in frequencies of different Fib-MET subpopulations (normalized for each population) across the groups: progressive SSc, stable SSc or controls. The Met^hi^_Fib are highlighted in red as significantly increased in progressive SSc. **C-D.** Frequencies of Met^hi^_Fib (C) or Met^low^_Fib (D) across the groups: progressive SSc, stable SSc or controls, shown as box plots. Statistical significance was determined by the Kruskal Wallis test with Dunn’s post-hoc test. 0.05>*P*>0.01*, 0.01>*P*>0.001**, *P*<0.001***. **E.** Representative images illustrating spatial distribution of the Fib-MET subsets across the groups: progressive SSc, stable SSc or controls, plotted on cell segmentation masks. HE images were included. This figure was created with Biorender.com. UMAP: Uniform Manifold Approximation and Projection; Fib: fibroblasts; Fib-MET: metabolically defined fibroblast subsets; SSc: systemic sclerosis; HE: hematoxylin and eosin.

### Metabolic changes in other cell types

We next performed unbiased clustering of endothelial cells (EC) based on metabolic markers and identified five distinct endothelial cells subsets, referred to as “metabolically-defined endothelial (End-MET) populations”: 1) Met^low^_EC, with a low metabolic activity; 2) TCA/OXPHOS^hi^PGC-1α^hi^_EC, with a high expression of TCA and OXPHOS enzymes and of PGC-1α; 3) Met^hi^_EC, with a high metabolic activity; 4) pmTOR^hi^CD98^hi^_EC, with a high activity of the mTOR pathway and an upregulation of the amino acid transporter CD98 and 5) Glycolysis^hi^TCA/OXPHOS^int^PPP^hi^_EC, with high levels of glycolysis enzymes, of the PPP enzyme G6PD and intermediate levels of the TCA/OXPHOS enzymes (Fig. S5A-C, E, F).

We next aimed to determine evidence of endothelial-to-mesenchymal transition (EndMT). We therefore evaluated the expression of αSMA across the End-MET populations. Met^hi^_EC subpopulation expressed the highest levels of αSMA as a prototypical marker of EndMT ^27^ (Fig. S5D). Thus, we provide evidence for a profound metabolic reprogramming in ECs undergoing EndMT in skin tissue.

As EndMT is thought to drive progression of skin fibrosis, we analyzed whether the frequencies of aSMA+ Met^hi^_EC are increased in SSc patients with progressive skin fibrosis (Fig. S6A). Indeed, the frequency of Met^hi^_EC is highly increased and that of the Met^low^_EC decreased in patients with progressive skin fibrosis compared with patients with stable skin fibrosis and with controls (Fig S6B-C). Furthermore, Glycolysis^hi^TCA/OXPHOS^hi^PPP^hi^_EC were numerically increased in patients with progressive skin fibrosis (Fig. S6B).

We also performed unbiased clustering of the macrophages (Mf) based on metabolic markers (Fig. S7A). We identified four “metabolically-defined macrophage (Mf-MET) populations”: 1) Met^hi^_Mf, with high metabolic activity; 2) Met^low^_Mf, with low metabolic activity; 3) Glycolysis^low^TCA/OXPHOS^hi^_Mf, with high expression of enzymes from the TCA/OXPHOS pathways, but low glycolytic activity and 4) Glycolysis^hi^TCA/OXPHOS^hi^_Mf, with high activity levels of both glycolysis and TCA/OXPHOS (Fig. S7B-E). The frequencies of the Met^hi^_Mf are highly increased, whereas the frequencies of Met^low^_Mf are numerically decreased in SSc patients with progressive skin fibrosis compared to patients with stable skin fibrosis and controls (Fig. S8).

Dermal cells such as fibroblasts, endothelial cells and macrophages in fibrotic skin thus show partial overlap in the metabolic phenotypes with Met^hi^ and Met^low^ in all cell types and TCA/OXPHOS^hi^PGC-1α^hi^ and pmTOR^hi^CD98^hi^ populations in fibroblasts and endothelial cells (Fig. 3, S5, S7). Changes in these metabolically defined populations tended to be homogenous, e.g. with consistent changes in e.g. Met^hi^ and Met^low^ populations of fibroblasts, endothelial cells and macrophages within individual patients and controls (Figure 4, S6, S8).

**Figure 5:**
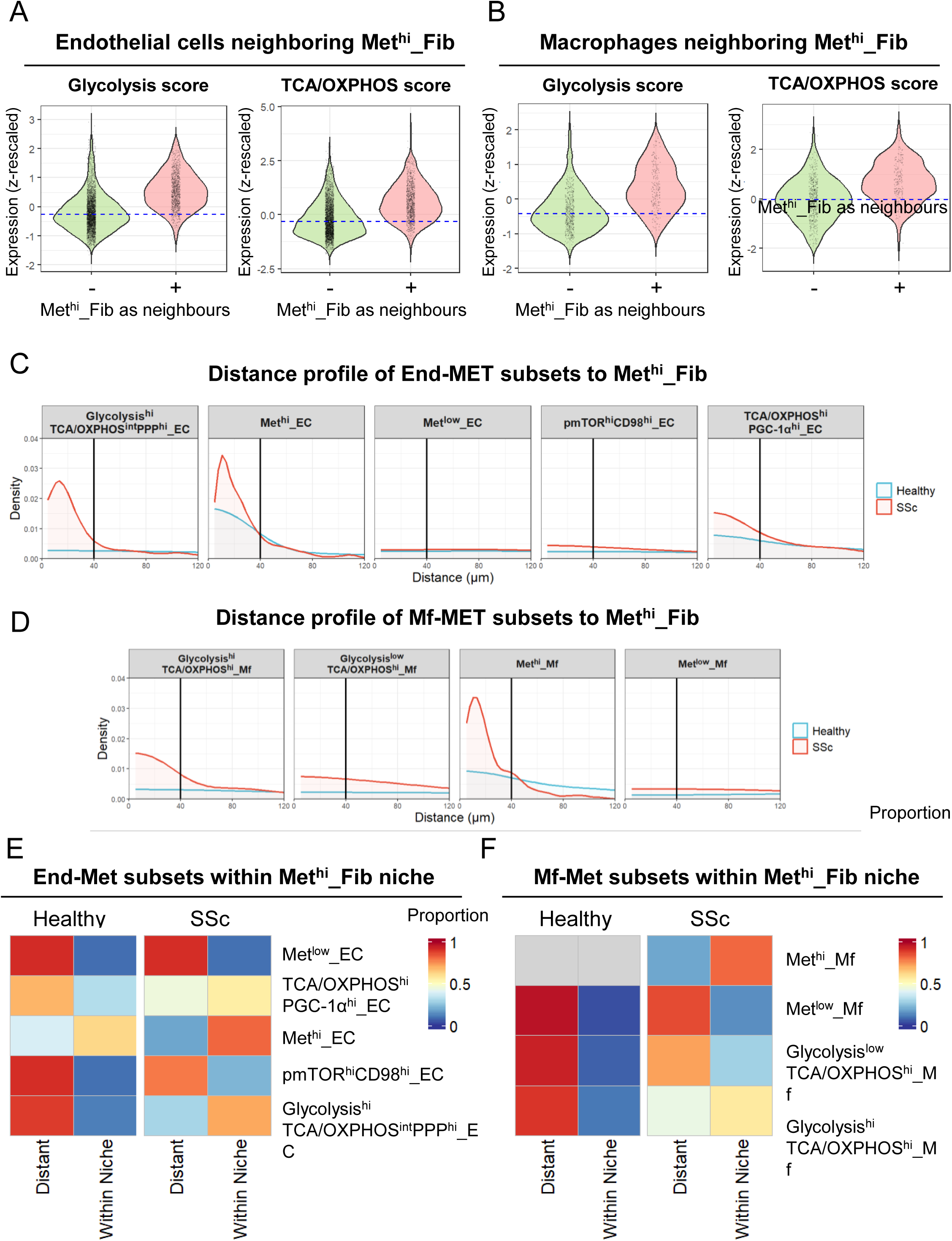
Spatial relationships of metabolically defined endothelial cell and macrophage subsets with Met^hi^_Fib A-B. Glycolysis score (calculated by averaging the expression of enzymes belonging to the glycolysis pathway: GLUT1, HK1, PFKL1, PKM2, LDHA, GAPDH), TCA/OXPHOS score (calculated by averaging the expression of enzymes belonging to the TCA cycle and OXPHOS: CS, OGDH, ATP5A, SDHA) in endothelial cells (A) or macrophages (B) that are neighboring Met^hi^_Fib. **C-D**. Distance of End-MET (C) or Mf-MET (D) subsets to Met^hi^_Fib in SSc patients and controls shown as density plots. **E-F.** Proportions of End-MET (E) or Mf-MET (F) subsets present within the Met^hi^_Fib niche or outside it in SSc patients or controls, shown as heatmaps. SSc: systemic sclerosis; Fib: fibroblasts; GLUT1: Glucose transporter 1; HK1: Hexokinase1; PFKL1: Phosphofructokinase Ligand 1; PKM2: Pyruvate kinase isoform M2; LDHA: Lactate dehydrogenase A; GAPDH: Glyceraldehyde 3-phosphate dehydrogenase; CS: Citrate synthase; OGDH: 2-oxoglutarate dehydrogenase; ATP5A: Mitochondrial adenosine triphosphatase alpha chain; SDHA: succinate dehydrogenase complex; End-MET: metabolically defined endothelial subsets; Mf-MET: metabolically defined macrophage subsets; TCA: tricarboxylic acid; OXPHOS: oxidative phosphorylation.

In contrast to the homogenous metabolic signatures in fibroblasts, endothelial cells and macrophages, the metabolic profile of keratinocytes is remarkably different. In contrast to all the other cell types, keratinocytes showed a prominent donor-to-donor variation, both in SSc patients and in controls (Fig. S9A, B). This higher inter-donor variation in the activity of metabolic pathways might reflect the direct exposure of keratinocytes to multiple environmental factors, in contrast to other cells located in deeper layers of the skin.

### Spatial determinants of the metabolic phenotypes of fibroblasts, endothelial cells and macrophages in SSc skin

We reasoned that the distinct metabolic profiles of fibroblasts might metabolically re-shape their local microenvironment and might induce changes in the metabolic profile of the neighboring cells. We observed that the glycolysis score and the TCA/OXPHOS score were enriched in EC and Mf that are neighboring the Met^hi^_Fib (Fig. 5A, B). We next analyzed the distribution of the distinct metabolic subsets of EC and Mf in the vicinity of Met^hi^_Fib. We observed a strong metabolic polarization in neighboring EC and Mf, with an enrichment in particular of Met^hi^_EC and of Met^hi^_Mf subsets in close spatial proximity to the Met^hi^_Fib (Fig. 5C, D). The enrichment of these Met^hi^_EC and Met^hi^_Mf subsets was particularly prominent in SSc (Fig. 5C, D). Furthermore, the Glycolysis^hi^TCA/OXPHOS^int^PPP^hi^_EC and the Glycolysis^hi^TCA/OXPHOS^hi^_Mf were enriched in the vicinity of Met^hi^_Fib only in SSc (Fig. 5C, D). Consistently, we observed that a high fraction of all Met^hi^_EC and Met^hi^_Mf were present within the microenvironment of Met^hi^_Fib (Fig. 5E, F). This fraction was low in controls but exceeded 80% in SSc patients (Fig. 5E, F). Glycolysis^hi^TCA/OXPHOS^int^PPP^hi^_EC and Glycolysis^hi^TCA/OXPHOS^hi^_Mf subsets were also enriched in the microenvironment of Met^hi^_Fib in SSc, with a fraction of at least 50% present within these niches (Fig. 5E, F).

In contrast, we observed only minor differences in the distribution of metabolically defined fibroblast subsets in the vicinity of vessels, of immune cell infiltrates, of the epidermis, or in the microenvironment of Met^hi^_EC or Met^hi^_Mf (Fig. S10, S11).

Taken together, these results highlight a strong spatial dependency of the metabolic polarization of EC and Mf on the metabolic profile of the fibroblasts, and suggest that, in particular in SSc, EC and Mf require spatial proximity to the Met^hi^_Fib to acquire a similar metabolic phenotype. In contrast, the metabolic profile of fibroblasts might be driven rather by cell intrinsic stimuli than by their local metabolic cues.

### Local microenvironment of the metabolically defined fibroblasts subsets in SSc skin

We next aimed to identify and characterize metabolic niches or spatial domains by clustering the distinct metabolic subsets of fibroblasts, endothelial cells and macrophages according to the composition of their cellular neighbors. We identified 10 distinct domains, or so-called cellular neighborhoods (CNs) (Fig. 6A). One of these identified the epidermis and was thus termed “Epi CN”. Another domain was mainly composed of immune cells and was termed “Immune CN”. A third domain was composed of two End-MET subsets and was termed “Vascular CN”. The remaining seven domains had a metabolically distinct fibroblast population in their composition, frequently among other metabolically defined EC or Mf subsets, and were thus termed Met-CN 1 to 7 (Fig. 6A). Several of these Met-CNs were composed of an End-MET or Mf-Met subtype with a similar metabolic profile to that of the Fib-MET subtype enriched in the same corresponding CN. Met-CN6, for example, was composed of Met^hi^_Fib, Met^hi^_EC and Met^hi^_Mf. Met-CN2 was composed of TCA/OXPHOS^hi^PGC-1α^hi^_Fib and TCA/OXPHOS^hi^PGC-1α^hi^_EC. Met-CN4 had in its composition Met^low^_Fib, Met^low^_EC and Met^low^_Mf (among other cell types) (Fig. 6A).

**Figure 6:**
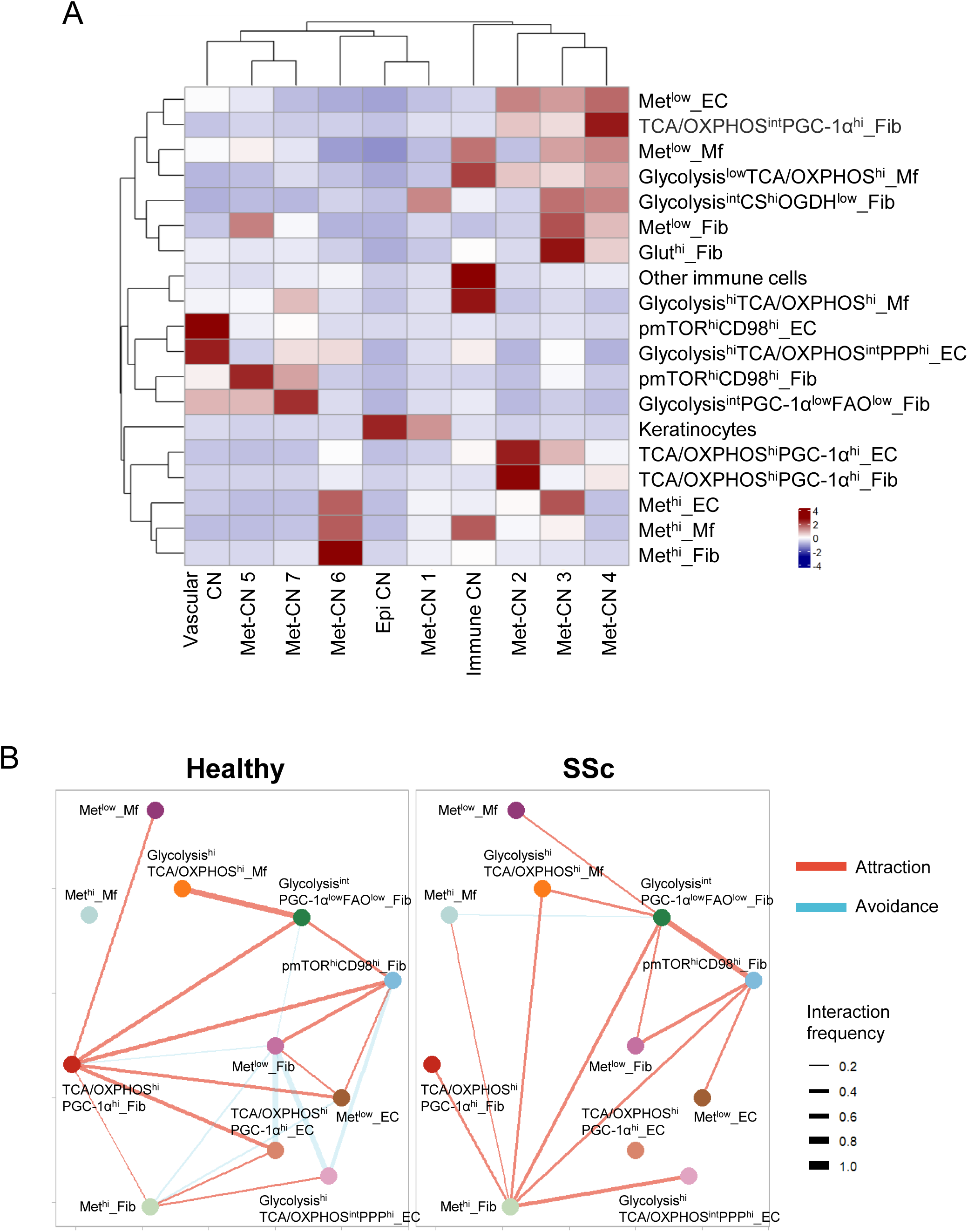
Metabolically defined cellular neighbourhoods and cell-cell interactions in SSc patients and controls. **A.** Cellular composition (including Fib-MET, End-MET and Mf-MET subsets) of metabolic CNs, shown as heatmap. The number of cells were normalised for each cell subset across all CNs. **B.** Network visualisation showing the interaction between indicated Fib-MET, End-MET and Mf-MET by pairwise interaction analysis, based on the cellular distribution and cell-cell spatial proximity in SSc skin and controls. The width of the lines represents the interaction frequencies, i.e. proportion of samples with statistically significant interactions, with red indicating attraction and blue indicating avoidance. SSc: systemic sclerosis; Fib-MET: metabolically defined fibroblast subsets; End-MET: metabolically defined endothelial subsets; Mf-MET: metabolically defined macrophage subsets.

**Figure 7:**
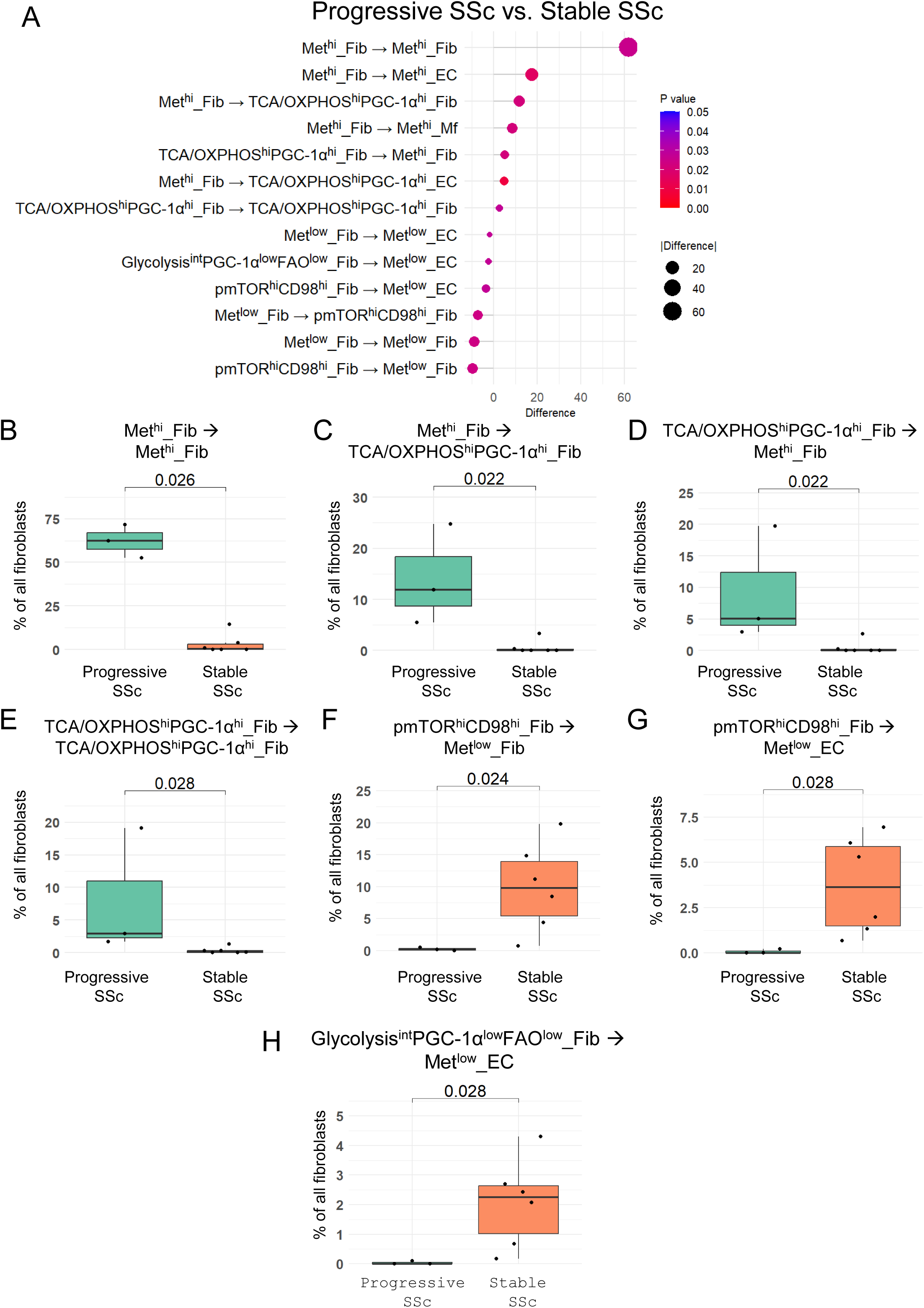
Changes in frequencies of metabolically defined fibroblast subsets in spatial proximity to distinct End-MET or Mf-MET subsets in progressive SSc. **A.** Dot plot illustrating the difference in frequencies of Fib-MET subsets in spatial proximity to distinct End-MET or Mf-MET subsets (among all fibroblasts) in progressive vs. stable SSc, shown for statistically significant comparisons. The size of the dots represents the module of the difference in frequency, and the color represents the p-value. **B-H.** Box plots illustrating changes in frequencies of Met^hi^_Fib in spatial proximity to other Met^hi^_Fib (B), or TCA/OXPHOS^hi^PGC-1α^hi^_Fib (C), or of TCA/OXPHOS^hi^PGC-1α^hi^_Fib in spatial proximity to Met^hi^_Fib (D) or to other TCA/OXPHOS^hi^PGC-1α^hi^_Fib (E), or of pmTOR^hi^CD98^hi^_Fib in spatial proximity to Met^low^_Fib (F) or Met^low^_EC (G), or of Glycolysis^int^PGC-1α^low^FAO^low^_Fib in spatial proximity to Met^low^_EC (H) (among all fibroblasts) in progressive vs. stable SSc. Statistical significance was determined by the Wilcoxon test. SSc: systemic sclerosis; Fib-MET: metabolically defined fibroblast subsets; End-MET: metabolically defined endothelial subsets; Mf-MET: metabolically defined macrophage subsets; Fib: fibroblasts, EC: endothelial cells; Mf: macrophages; TCA: tricarboxylic acid; OXPHOS: oxidative phosphorylation; FAO: fatty acid oxidation.

Furthermore, the Met-CNs demonstrated distinct expression patterns of the metabolic markers, with Met-CN6 having the highest expression of most metabolic markers, Met-CN4 having among the lowest expression of these markers, and Met-CN2 having high expression of the enzymes from TCA cycle and OXPHOS, as well as of PGC-1α, with a lower expression of glycolytic enzymes (Fig. S12A). These findings suggest a spatial aggregation of endothelial cells and macrophages to fibroblasts with a similar metabolic profile.

We further compared the changes in frequencies of Met-CN in SSc patients compared to controls. Met-CN6 was significantly more frequent in SSc donors, and it was not present in most of the controls (Fig. S12B). All donors with a high frequency of Met-CN6 (more than 25% of all Met-CNs) were patients with progression of skin fibrosis (Fig. S12C).

The composition of the Met-CN demonstrated important differences in SSc patients compared to controls. While the main Fib-MET subset in the composition of each Met-CN was generally preserved, several Fib-MET subsets were enriched in a distinct Met-CN in SSc compared to controls. Met^hi^_Fib, for example, were enriched in Met-CN2 in controls, and in Met-CN6 in SSc (Fig. S13A). Similarly, Met^hi^_EC and Met^hi^_Mf were enriched in Met-CN2 in controls and in Met-CN6 in SSc (Fig. S13B, C), highlighting that, in controls, these metabolic cell profiles do not form a distinct metabolic niche, but rather assemble in niches formed by cells with different metabolic phenotypes.

We next performed cell-cell interaction analysis to characterize changes in the interaction partners of the metabolically defined fibroblasts, EC and Mf populations in SSc. We observed a remarkably different interaction network in the skin of SSc patients compared to controls (Fig. 6B, S14). The Met^hi^_Fib in SSc demonstrated several interactions that are present only or are highly enriched in SSc, i.p. with Glycolysis^hi^TCA/OXPHOS^int^PPP^hi^_EC, Met^hi^_Mf, Glycolysis^hi^TCA/OXPHOS^hi^_Mf, pmTOR^hi^CD98^hi^_Fib, Glycolysis^int^PGC-1α^low^FAO^low^_Fib and TCA/OXPHOS^hi^PGC-1α^hi^_Fib (Fig. 6B, S14).

Of note, other fibroblast populations also demonstrated major changes in their cellular interactions, despite only modest changes in frequency in SSc. The TCA/OXPHOS^hi^PGC-1α^hi^_Fib population loses most of its interactions in SSc (Fig. 6B, S14). The Glycolysis^int^PGC-1α^low^FAO^low^_Fib subset interacts specifically with Met^low^_Mf and Met^low^_Fib, and upregulates its interactions with pmTOR^hi^CD98^hi^_Fib in SSc (Fig. 6B, S14).

### Correlations of metabolically distinct fibroblast subsets and their cellular interactions with clinical features of SSc

We next aimed to evaluate the association between frequencies of metabolically defined fibroblast subsets and clinical features in SSc. Since the Fib-MET subsets had different interaction partners in SSc donors compared to controls, we reasoned that incorporating the spatial relationships between the distinct metabolically defined cellular subpopulations can refine this analysis, as variations in cell-cell communication may be differentially associated with clinical outcomes.

We first evaluated associations between progression of skin fibrosis and the frequency of each Fib-MET subpopulations interacting with metabolically defined fibroblasts, endothelial cells or macrophage populations (Fig. 7A). The frequency of Met^hi^_Fib in spatial proximity to other Met^hi^_Fib, to Met^hi^_EC, to Met^hi^_Mf, or to TCA/OXPHOS^hi^PGC1α^hi^_Fib or TCA/OXPHOS^hi^PGC1α^hi^_EC was strongly increased in SSc patients with progressive skin fibrosis (Fig. 7A-C). The shifts in frequencies of Met^hi^_Fib interacting with other metabolically defined cell subsets were milder and/or did not reach statistical significance (Fig. S15). The frequency of the TCA/OXPHOS^hi^PGC-1α^hi^_Fib interacting with other TCA/OXPHOS^hi^PGC-1α^hi^_Fib or with Met^hi^_Fib was also higher in patients with progressive skin fibrosis (Fig. 7A, D, E).

In contrast, several interactions pairs involving Met^low^ such as mTOR^hi^CD98^hi^_Fib interacting with Met^low^_Fib or Met^low^_EC, or Glycolysis^int^PGC-1α^low^FAO^low^_Fib interacting with Met^low^_EC, were enriched in SSc patients with stable skin fibrosis (Fig. 7A, F-H).

We next evaluated associations between skin fibrosis severity, as measured by mRSS, and the frequency of each Fib-MET subset interacting with metabolically defined fibroblasts, endothelial cells or macrophage populations (Fig. 8A). The frequency of Met^hi^_Fib in spatial proximity to pmTOR^hi^CD98^hi^_Fib, Met^low^_Fib, pmTOR^hi^CD98^hi^_EC, Glycolysis^hi^TCA/OXPHOS^int^PPP^hi^_EC or Glycolysis^hi^TCA/OXPHOS^hi^_Mf correlated with the mRSS (Fig. 8A-E). Furthermore, the frequencies of pmTOR^hi^CD98^hi^_Fib interacting with Met^hi^_Fib, as well as the frequencies of Glycolysis^int^PGC-1α^low^FAO^low^_Fib interacting with Met^hi^_Fib or Met^hi^_Mf correlated with the mRSS (Fig. 8A, F-H). These interactions correlated specifically with mRSS and not with other demographic or clinical features such as age, disease duration or CRP (Fig. S16). Of note, the frequency of Met^hi^_Fib, pmTOR^hi^CD98^hi^_Fib or Glycolysis^int^PGC-1α^low^FAO^low^_Fib demonstrated a much weaker correlation with the mRSS, when not accounting for their interaction partners (Fig. S17), or when they interacted with other metabolically defined cell subsets than those mentioned above (Fig. S18). Furthermore, the interactions that correlate to mRss are distinct from the interactions enriched in progressive SSc (Fig. 7, Fig. 8 and Fig. S18). This suggests that specific metabolic cell-cell communication networks might be involved in progression of fibrotic skin remodeling and in extensive skin fibrosis in SSc.

**Figure 8:**
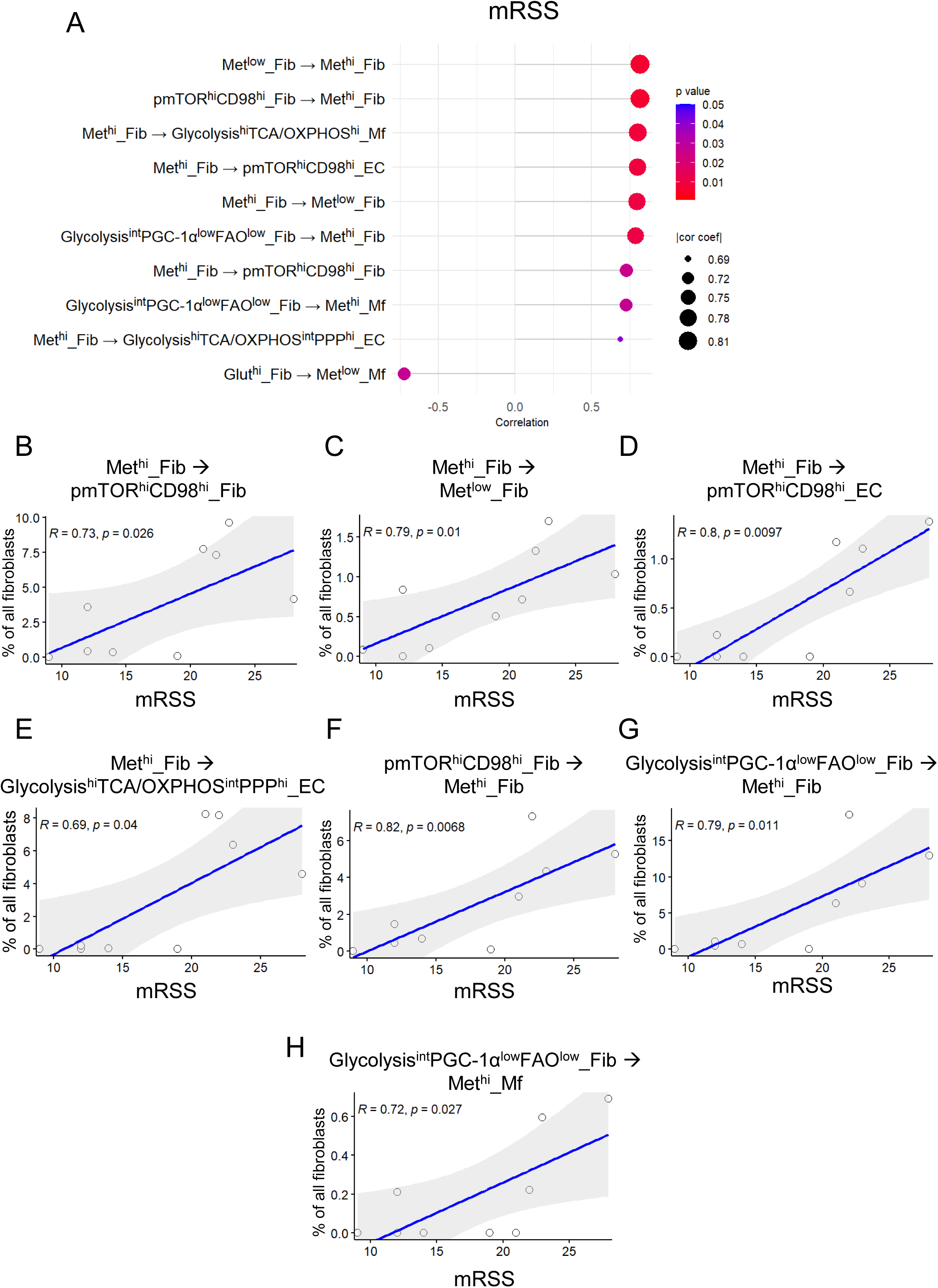
Correlations of mRSS with the frequencies of metabolically defined fibroblast subsets in spatial proximity to distinct End-MET or Mf-MET subsets in SSc. **A.** Dot plot illustrating the correlation of mRSS with the frequency of Fib-MET subsets in spatial proximity to distinct End-MET or Mf-MET subsets (among all fibroblasts), shown for statistically significant comparisons. The size of the dots represents the coefficient of correlation, and the color represents the p-value. **B-H.** Scatter plots illustrating the correlation between mRSS and the frequency of Met^hi^_Fib in spatial proximity to pmTOR^hi^CD98^hi^_Fib (B), Met^low^_Fib (C), pmTOR^hi^CD98^hi^_EC (D), Glycolysis^hi^TCA/OXPHOS^int^PPP^hi^_EC (E), or of pmTOR^hi^CD98^hi^_Fib in spatial proximity to Met^hi^_Fib (F), or of Glycolysis^int^PGC-1α^low^FAO^low^_Fib in spatial proximity to Met^hi^_Fib (G) or Met^hi^_Mf (H) (among all fibroblasts). Statistical significance was determined by Spearman’s correlation analysis. Linear regression lines with the 95% CI, the Spearman’s rank correlation coefficient and the p-value are included in the scatter plots. SSc: systemic sclerosis; Fib-MET: metabolically defined fibroblast subsets; End-MET: metabolically defined endothelial subsets; Mf-MET: metabolically defined macrophage subsets; Fib: fibroblasts, EC: endothelial cells; Mf: macrophages; mRSS: modified Rodnan skin score; TCA: tricarboxylic acid; OXPHOS: oxidative phosphorylation; FAO: fatty acid oxidation.

## Discussion

Our study provides the first spatially resolved metabolic profiling of human skin from SSc patients and normal individuals at single-cell resolution. IMC with an antibody panel targeting a broad spectrum of metabolic enzymes and regulators that has previously been shown to correlate well with the results of metabolic flux assays ^18^, provided us with the unique opportunity to simultaneously assay several key metabolic pathways. We identify a specific metabolic profile in the dermal compartment that is associated with progression of skin fibrosis in SSc patients. While the metabolic profile of SSc patients with stable or regressive skin fibrosis did not show differences to normal skin of controls, SSc patients with progressive skin fibrosis showed remarkable changes in multiple metabolic pathways with most pronounced changes observed for the TCA cycle, oxidative phosphorylation, glycolysis and ROS production. Of note, these metabolic changes were only observed in patients with progressive fibrosis, whereas other clinical parameters such as extent of skin fibrosis, fibrosis of other organs or disease duration were not associated with metabolic changes compared to normal skin. The findings of upregulated HIF1α and NOX4 confirmed the validity of our IMC-based metabolic assessments, as tissue hypoxia and enhanced ROS production have previously been reported by other methods ^28,29^. These findings provide evidence that metabolic activation with enhanced oxidative phosphorylation and glycolysis is required to promote fibrotic tissue remodeling. Moreover, this metabolic activation pattern may serve as a surrogate marker for active fibrotic remodeling. However, further studies on additional patients and cohorts will be required to confirm these findings for the skin and evaluate whether metabolic activation is also associated with progressive fibrotic remodeling in other organs.

The IMC approach also enabled us to define subpopulations of fibroblasts, endothelial cells and macrophages based on the metabolic phenotype of each individual cell. Previous studies demonstrated that fibroblasts and endothelial cells are heterogeneous at a transcriptome or protein expression levels in SSc and other fibrotic diseases ^9,20,26,30–32^; however, the metabolic heterogeneity of fibroblasts or endothelial cells not been studied beyond the transcriptional level thus far. Using the metabolism-based clustering approach, we identified eight metabolically distinct subpopulations of fibroblasts, five subpopulations of endothelial cells and four subpopulations of macrophages, distinct with regards to their expression levels of key enzymes involved in metabolic pathways such as glycolysis, TCA/OXPHOS, FAO, pentose phosphate pathway or amino acid transport, and thus with regards to the activity of these pathways.

Progressive skin fibrosis was associated with major shifts in metabolically defined subpopulations of fibroblasts, endothelial cells and macrophages with shifts from metabolically inactive phenotypes towards metabolically highly active subpopulations, with high glycolytic and TCA/OXPHOS activity in all three lineages. Of note, these metabolically highly active populations also upregulate HIF1α and NOX4, suggesting that they are present in regions of hypoxia and with high ROS content. In contrast, other metabolically defined fibroblast populations, i.e. pmTOR^hi^CD98^hi^, Glut^hi^ and Glycolysis^int^PGC1-α^low^FAO^low^ fibroblasts, were numerically increased in SSc patients with stable or regressive skin fibrosis compared to patients with progressive skin fibrosis or controls.

For fibroblasts, we demonstrate that these shifts in metabolically defined subpopulations are directly implicated into the pathogenesis of fibrotic tissue remodeling, as they promote transition from resting fibroblasts into myofibroblasts. Met^hi^_Fib express the highest levels of the prototypical myofibroblast marker αSMA, but also of other activation markers such as Cdh11 and FAP. This finding is consistent with the observation that myofibroblasts upregulate glycolysis, as described by ex vivo flux assays ^33,34^. The Met^hi^_Fib have the highest levels of TCA/OXPHOS, high levels of PGC-1α and TFAM, as well as high levels of CPT1A and high FAO activity. This contrasts with previous reports, which demonstrated reduced FAO activity of fibroblasts in tissue fibrosis ^35–37^. However, previous studies could not capture the metabolic fibroblast heterogeneity directly in fibrotic tissue. We show that other fibroblast subsets such as the Glycolysis^int^PGC1-α^low^FAO^low^_Fib resemble the phenotype described in previous reports, with intermediate level of glycolysis activity and low levels of FAO. These fibroblasts may account for the previously described phenotype observed in cell culture experiments.

Furthermore, metabolic reprogramming with upregulation of glycolysis and TCA/OXPHOS, as well as of hypoxia and ROS signaling was also observed in endothelial cells undergoing EndMT. Together, these findings suggest that activation of glycolysis and TCA/OXPHOS, accompanied by upregulation of hypoxia and ROS signaling, is a common prerequisite for transition of mesenchymal cells towards a profibrotic myofibroblast phenotype that might drive progression of skin fibrosis.

The Met^hi^_Mf upregulate both glycolysis and TCA/OXPHOS and thus show a metabolic phenotype with characteristics of both M1 and M2 polarized macrophages. Monocyte/macrophages that express phenotypic markers of both M1 and M2 have been previously described in SSc ^38^. However, it was not known whether this is indeed due to the presence of an intermediate macrophage polarization state, or rather due to the aberrant expression of M1 markers on macrophages with an M2 functional polarization.

Characterization of the local microenvironment of metabolically defined subpopulations revealed that the metabolic alterations occur particularly in specific cellular niches, rather than in histologic compartments. These niches seem to demonstrate a metabolic hierarchy. Several lines of evidence indicate that fibroblasts metabolically shape their local microenvironment and modulate the metabolism in surrounding cells to form specific metabolic niches in SSc skin. The microenvironment of Met^hi^_Fib demonstrates profound accumulation of metabolically active endothelial cells and leukocytes, whereas Met^hi^_Fib are locally only mildly enriched near Met^hi^_ECs or Met^hi^_Mf. Moreover, several metabolically defined cellular neighborhoods are predominantly defined by a particular fibroblast subpopulation. The endothelial cells and/or macrophages show a similar metabolic phenotype to the fibroblast population present in the respective neighborhood. Furthermore, even if in controls the Met^hi^_Fib population does not form its own neighborhood, but it is rather present in the neighborhood of another fibroblast subset, the metabolically high endothelial cells and macrophages are present in the same neighborhood, highlighting a strong spatial inter-dependency.

These metabolic niches may create a permissive microenvironment that facilitates cellular interactions and promotes profibrotic phenotypes of the individual cell types. Although functional studies are required to ultimately confirm this hypothesis, it is supported by profound increase in cellular interactions of metabolically active fibroblasts in the skin of SSc patients with several other subsets of fibroblasts, endothelial cells or macrophages that have intermediate-high levels of glycolysis or TCA/OXPHOS activity. In line with this, *ex vivo* experiments demonstrated that SSc fibroblasts acidify their environment due to their high glycolytic activity, which promotes EndMT in endothelial cells ^39^. Furthermore, profibrotic cytokines released by macrophages, such as TGF-β and IL-13 or stabilization of HIF-1α by tissue hypoxia can promote glycolysis in fibroblasts and other cells within their niche ^12^.

The interactions between metabolically distinct dermal cells are directly relevant for clinical outcomes. The frequencies of metabolically defined fibroblast subsets were changed in patients with progression of skin fibrosis, when these subsets were interacting with particular cell populations. Furthermore, while the frequencies of fibroblast subsets did not correlate with the extent of skin fibrosis when not accounting for their interactions, we identified strong and specific correlations with mRSS, but not with other clinical features, of the frequencies of Met^hi^_Fib, of pmTOR^hi^CD98^hi^_Fib or Glycolysis^int^PGC-1α^low^FAO^low^_Fib, when they were in spatial proximity of particular metabolically defined cell types. These associations with the extent or with progression of skin fibrosis were much weaker when these cell populations had different interaction partners, providing further evidence that specific metabolic niches might facilitate fibrotic skin remodeling in SSc.

In summary, using a single cell, spatially resolved metabolic phenotyping approach, we identified a distinct metabolically active profile with high activity of glycolysis, TCA/OXPHOS, hypoxia and ROS signaling in fibroblasts, endothelial cells and macrophages of patients with progressive skin fibrosis. These metabolically active profiles were associated with expression of profibrotic activation markers in fibroblasts and endothelial cells. We further characterized metabolic niches in SSc skin and their hierarchical organization, where metabolically active fibroblasts might induce a similar metabolic phenotype of endothelial cells and macrophages, facilitating profibrotic cell-cell communication networks. Consistently, specific interactions between metabolically defined cell populations are strongly associated with the extent or progression of skin fibrosis. These results provide a rationale for therapeutic targeting of specific metabolically defined cell subsets and their interactions.

## Author contributions

JHWD and AEM designed the study. VD, YNL, TF, ARR, AHG, BST, RN, ML and AEM were involved in acquisition and analysis of data. VD, YNL, TF, AHG, ML, CB, GS, JHWD and AEM were involved in interpretation of data. ML, CB and GS provided essential samples. All authors were involved in manuscript preparation and proof-reading.

## Acknowledgements

We thank Christoph Liebel, Philipp Steinbrecher and Lukas Sokolowski for excellent technical assistance. The project was supported by the following grants: “Großgeräteantrag” of the German Research Foundation (DFG), INST 901095-1 FUGG / AOBJ 659788 / ID 424726560, Grants DI 1537/17-1, DI 1537/20-1, DI 1537/22-1, DI 1537/23-1, DI 1537/27-1, DI 1537/28-1 of the German Research Foundation (JHWD), MA 9219/2-1 of the German Research Foundation (AEM), SFB CRC1181 (project C01) of the German Research Foundation, grants 2021_EKEA.03 (AEM) and 2022_EKMS.02 (AEM) of the Else-Kröner-Fresenius-Foundation, The Edith Busch and World Scleroderma Foundation Research Grant Programme 2022-2023 (AEM), the German Federal Ministry of Education and Research (BMBF), MASCARA program, TP 2 (01EC1903A), the Research Committee of the Medical Faculty of the Heinrich-Heine University Düsseldorf (Forschungskommission; ID 2022-18, ID 2023-33 and ID 2023-31 to AHG, AEM and JHWD, respectively), an unrestricted research grant from the Hiller-Foundation (JHWD) and a Career Support Award of Medicine of the Ernst Jung Foundation (JHWD). Our analysis was supported by a de.NBI Cloud project (YNL) within the German Network for Bioinformatics Infrastructure (de.NBI) and ELIXIR-DE (Forschungszentrum Jülich and W-de.NBI-001, W-de.NBI-004, W-de.NBI-008, W-de.NBI-010, W-de.NBI-013, W-de.NBI-014, W-de.NBI-016, W-de.NBI-022).

---

**Figure S1:**
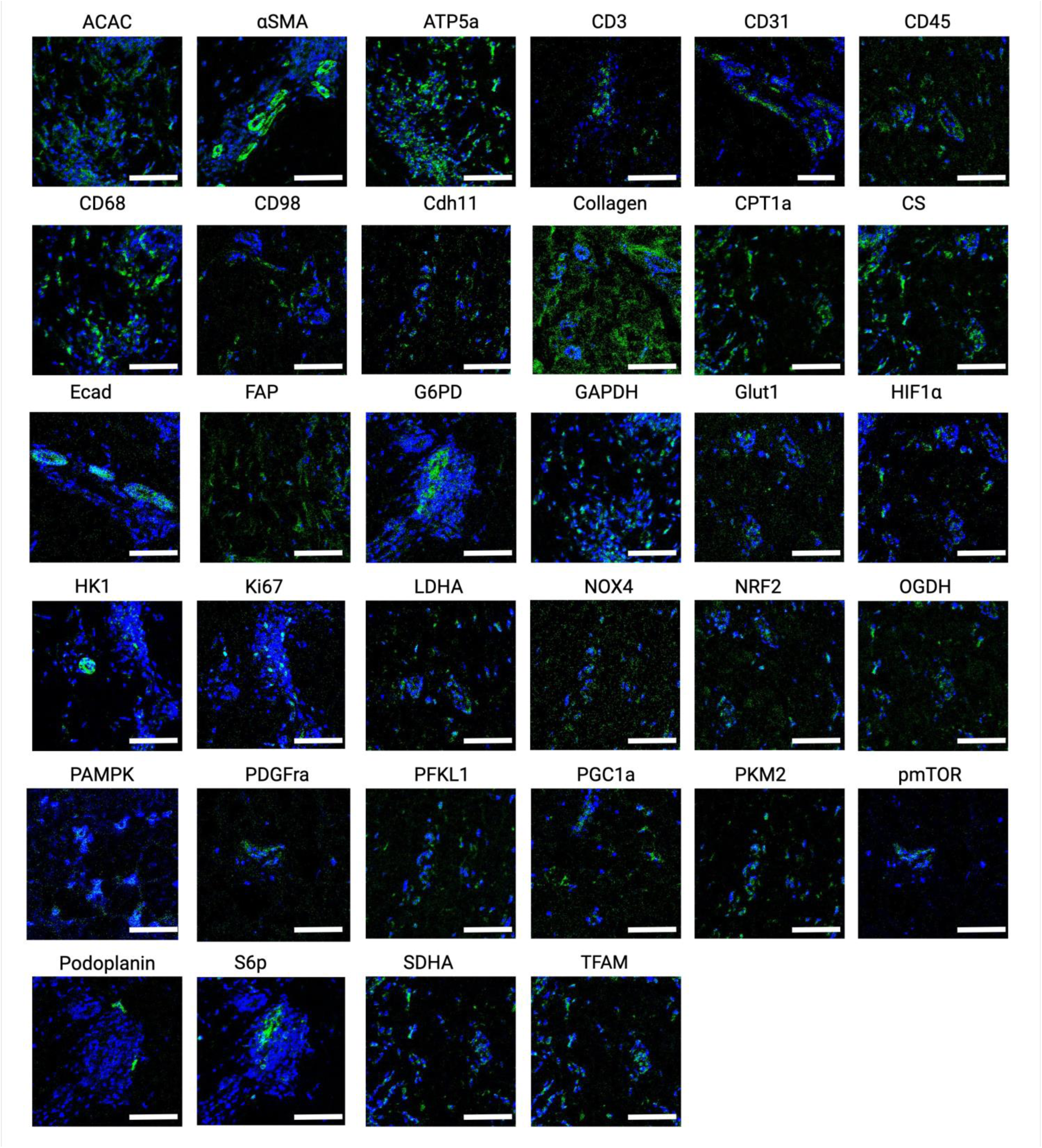
Spatial distribution of marker expression included in the IMC panel. Representative images illustrating spatial distribution of the signals of ACAC, αSMA, ATP5A, CD3, CD31, CD45, CD68, CD98, Cadherin 11, Collagen type 1, CPT1A, CS, E-Cadherin, FAP, G6PD, GAPDH, GLUT1, HIF1α, HK1, KI67, LDHA, NOX4, NRF2, OGDH, pAMPK, PDGFRα, PFKL1, PGC1α, PKM2, pmTOR, Podoplanin, pS6, SDHA and TFAM at pixel level. Scale bars: 75µm. This figure was created with Biorender.com. IMC: imaging mass cytometry.

**Figure S2:**
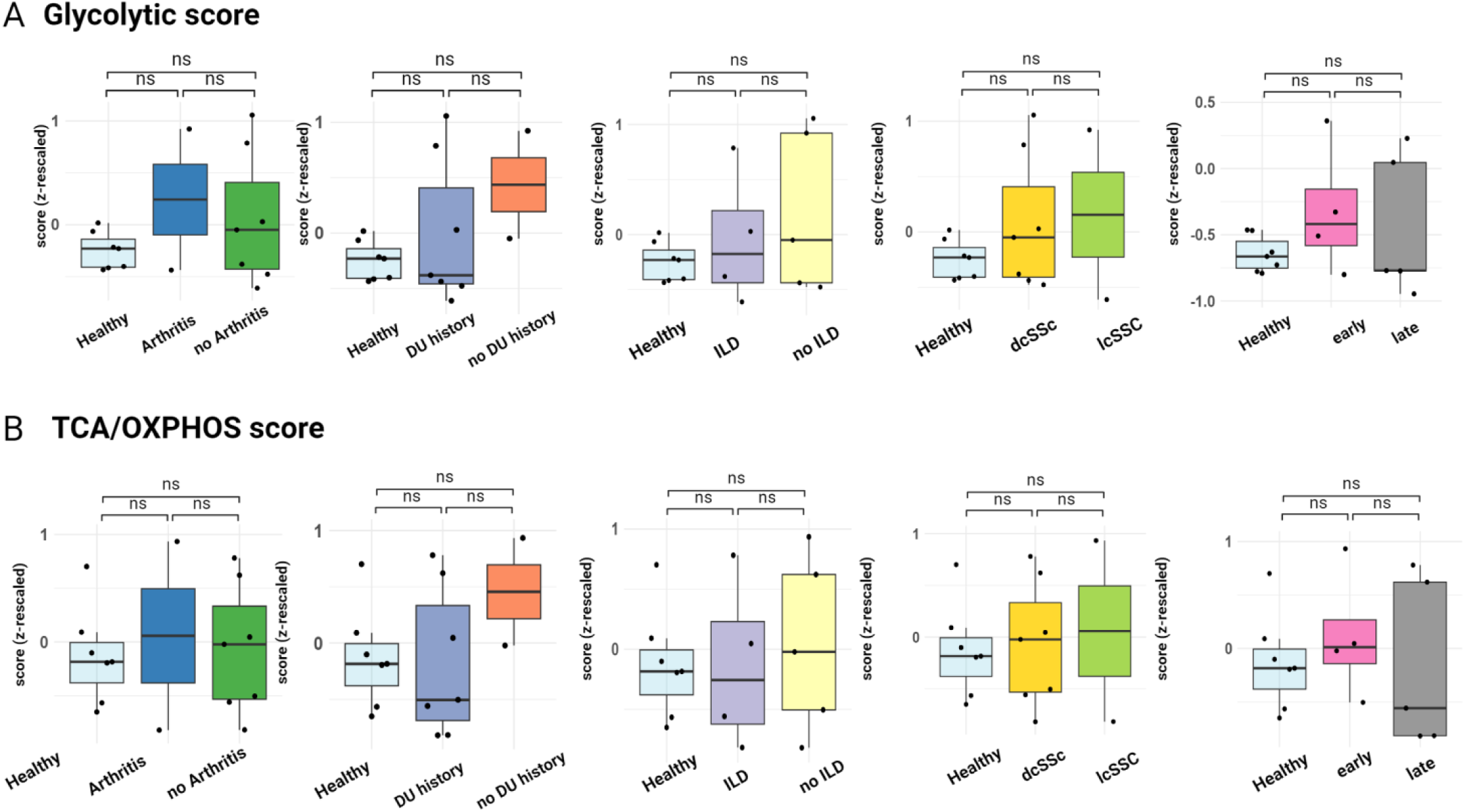
The glycolysis and the TCA/OXPHOS score are not associated with other clinical features of SSc except progression of skin fibrosis A-B. Glycolysis score (A) (calculated by averaging the expression of enzymes belonging to the glycolysis pathway: GLUT1, HK1, PFKL1, PKM2, LDHA, GAPDH) and TCA/OXPHOS score (B) (calculated by averaging the expression of enzymes belonging to the TCA cycle and OXPHOS: CS, OGDH, ATP5A, SDHA) in dermal cells across the groups: presence of arthritis, history of digital ulcers, presence of interstitial lung disease, diffuse or limited SSc, early or late SSc, shown as box plots. This figure was created with Biorender.com. SSc: systemic sclerosis; GLUT1: Glucose transporter 1; HK1: Hexokinase1; PFKL1: Phosphofructokinase Ligand 1; PKM2: Pyruvate kinase isoform M2; LDHA: Lactate dehydrogenase A; GAPDH: Glyceraldehyde 3-phosphate dehydrogenase; CS: Citrate synthase; OGDH: 2-oxoglutarate dehydrogenase; ATP5A: Mitochondrial adenosine triphosphatase alpha chain; SDHA: succinate dehydrogenase complex; TCA: tricarboxylic acid; OXPHOS: oxidative phosphorylation.

**Figure S3:**
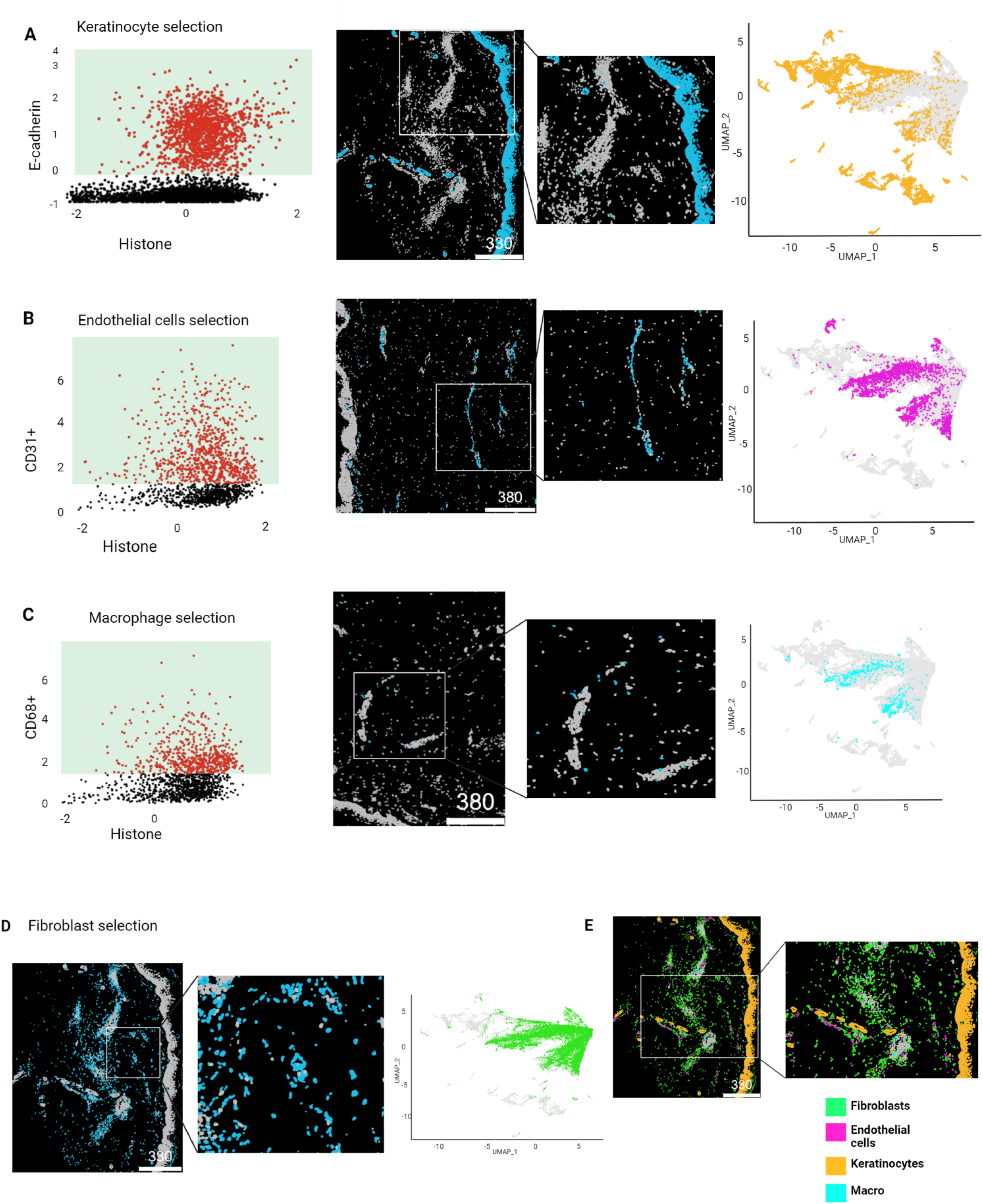
Gating strategy for identification of keratinocytes, endothelial cells, macrophages and fibroblasts A-D. Scatter plots illustrating the expression of E-Cadherin (A), CD31 (B) or CD68 (C), including rectangles that illustrate selection of positive cells for E-Cadherin, i.e. keratinocytes (A), for CD31, i.e. endothelial cells (B) and for CD68 from CD45+ cells, i.e. macrophages (C), with cells negative for E-Cadherin, CD31 and CD45 as fibroblasts (D). Representative images illustrating spatial distribution of keratinocytes (A), endothelial cells (B), macrophages (C) and fibroblasts (D) plotted on cell segmentation masks are included. UMAP plots generated based on expression of metabolic markers and colour coded by the identity of the main populations: keratinocytes (A), endothelial cells (B), macrophages (C) and fibroblasts (D) are also included. **E:** Representative images illustrating spatial distribution of keratinocytes, endothelial cells, macrophages and fibroblasts plotted on cell segmentation masks. This figure was created with Biorender.com. E-cad: E-cadherin; UMAP: Uniform Manifold Approximation and Projection.

**Figure S4:**
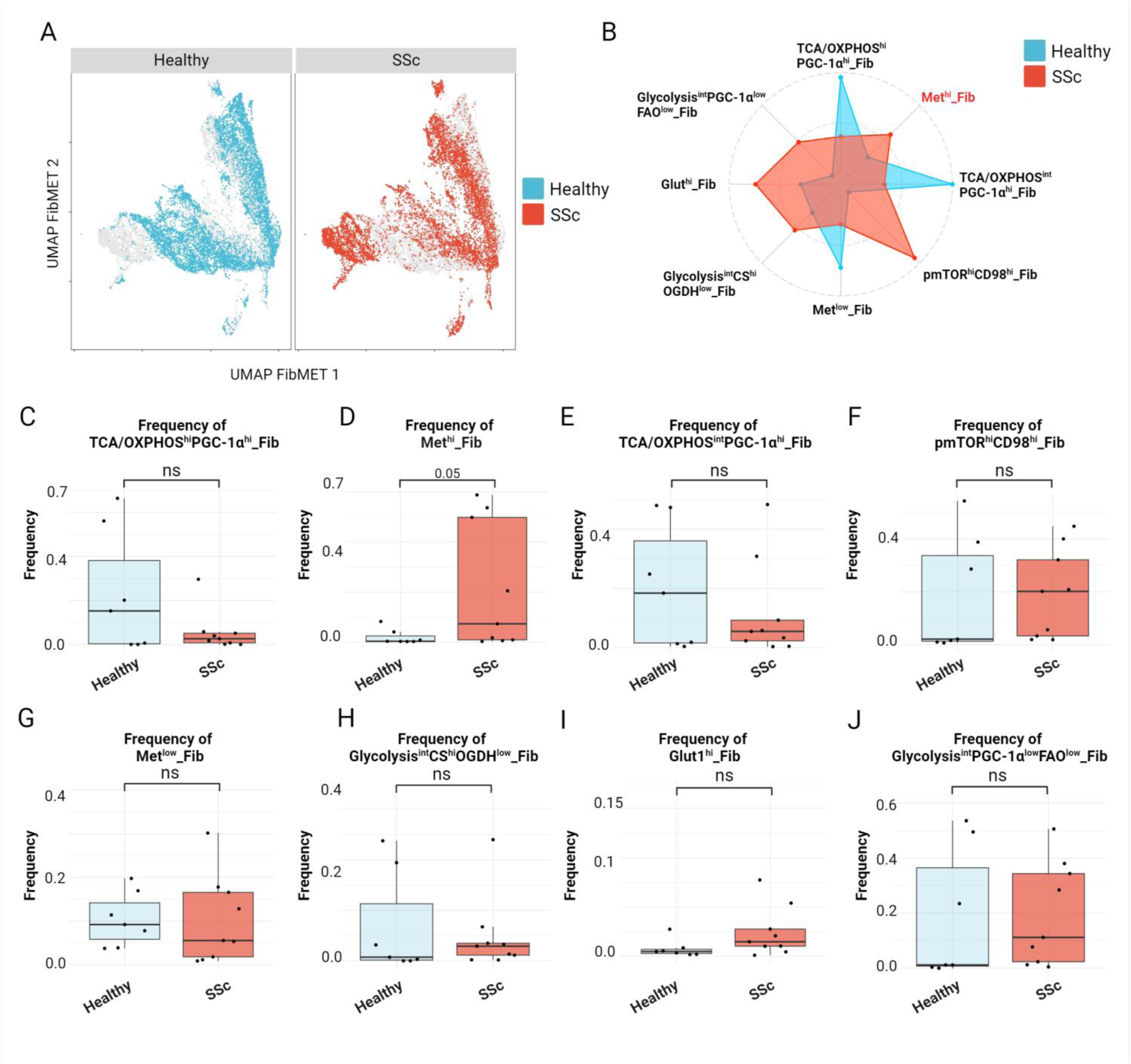
Shifts in frequencies of metabolically defined fibroblast subpopulations in SSc patients. **A.** UMAP plots generated based on expression of metabolic markers and colour coded by the groups: SSc or controls. **B.** Radar chart illustrating the shifts in frequencies of different Fib-MET subpopulations (normalized for each population) across the groups: SSc or controls. The Met^hi^_Fib are highlighted in red. **C-J.** Frequencies of TCA/OXPHOS^hi^PGC-1α^hi^_Fib (C), Met^hi^_Fib (D), TCA/OXPHOS^int^PGC-1α^hi^_Fib (E), pmTOR^hi^CD98^hi^_Fib (F), Met^low^_Fib (G), Glycolysis^int^CS^hi^OGDH^low^_Fib (H), Glut1^hi^_Fib (I) and Glycolysis^int^PGC-1α^low^FAO^low^_Fib across the groups: SSc or controls, shown as box plots. Statistical significance was determined by the Wilcoxon test. 0.05>*P*>0.01*, 0.01>*P*>0.001**, *P*<0.001***. This figure was created with Biorender.com. UMAP: Uniform Manifold Approximation and Projection; Fib: fibroblasts; Fib-MET: metabolically defined fibroblast subsets; SSc: systemic sclerosis.

**Figure S5:**
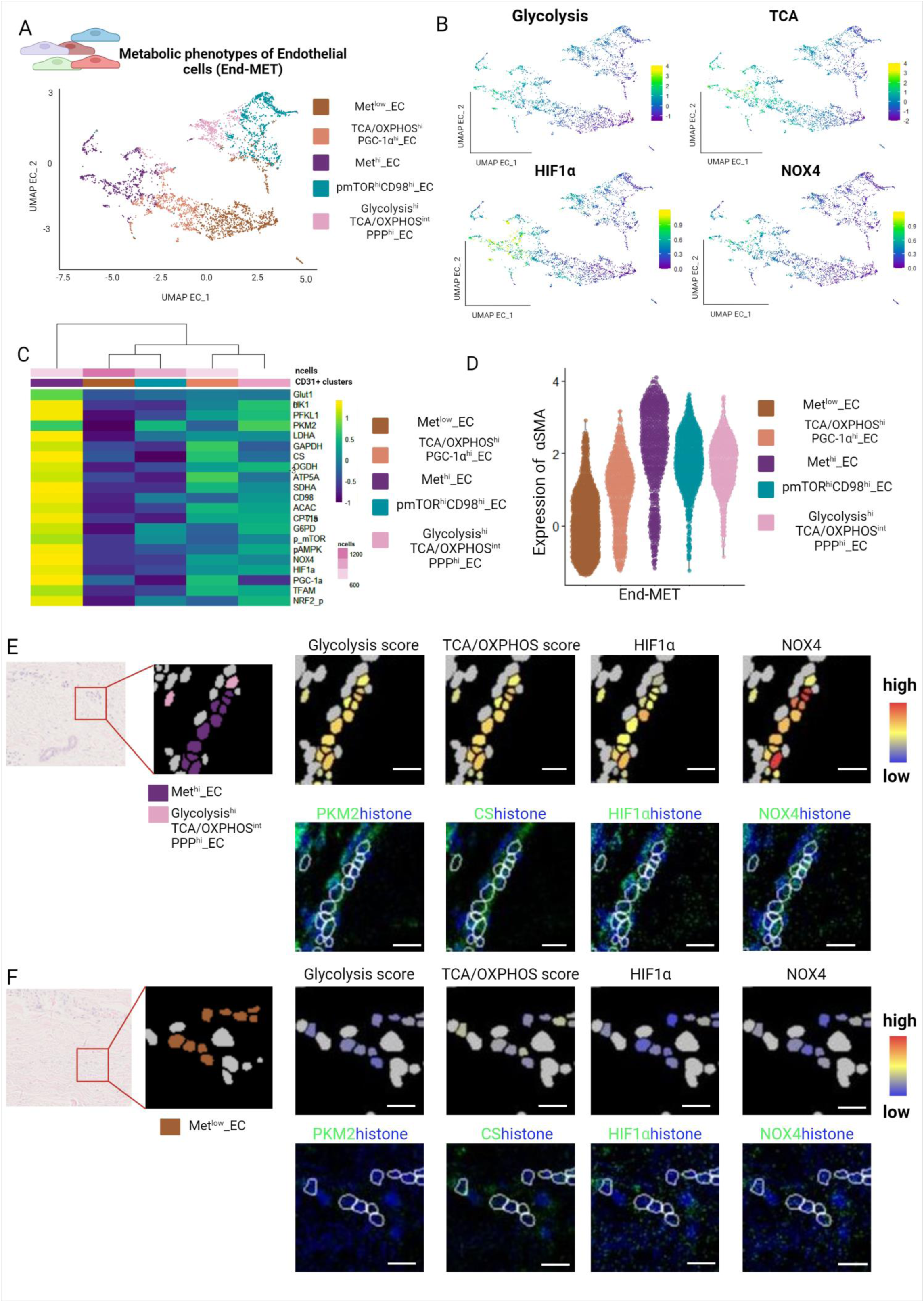
Identification of metabolically defined populations of endothelial cells. **A.** UMAP plots generated based on expression of metabolic markers and colour coded by metabolically defined endothelial cell subsets, and illustration of the distinct endothelial cell subsets. **B.** Glycolysis score (calculated by averaging the expression of enzymes belonging to the glycolysis pathway: GLUT1, HK1, PFKL1, PKM2, LDHA, GAPDH), TCA/OXPHOS score (calculated by averaging the expression of enzymes belonging to the TCA cycle and OXPHOS: CS, OGDH, ATP5A, SDHA), hypoxia (expression of HIF1α) and ROS (expression of NOX4) signaling in endothelial cells plotted on UMAP. **C.** Expression of metabolic markers across metabolically defined endothelial cell subsets, plotted as heatmap. **D.** Expression of αSMA across metabolically defined endothelial cell subsets, shown as violin plot. **E-F.** Representative images illustrating spatial distribution of the Met^hi^_EC, Glycolysis^hi^TCA/OXPHOS^int^PPP^hi^_EC (E) and of Met^low^_EC (F), along glycolysis and TCA/ETC scores and hypoxia and ROS signaling, plotted on cell segmentation masks, and images showing expression of PKM2 (a glycolytic enzyme), CS (an enzyme from the TCA cycle), HIF1α (hypoxia signaling) and NOX4 (ROS signaling) at pixel level. Corresponding HE images were included. Scale bars: 25 µm. This figure was created with Biorender.com. UMAP: Uniform Manifold Approximation and Projection; HE; hematoxylin and eosin; EC: endothelial cells; TCA: tricarboxylic acid; OXPHOS: oxidative phosphorylation; GLUT1: Glucose transporter 1; HK1: Hexokinase1; PFKL1: Phosphofructokinase Ligand 1; PKM2: Pyruvate kinase isoform M2; LDHA: Lactate dehydrogenase A; GAPDH: Glyceraldehyde 3-phosphate dehydrogenase; CS: Citrate synthase; OGDH: 2-oxoglutarate dehydrogenase; ATP5A: Mitochondrial adenosine triphosphatase alpha chain; SDHA: succinate dehydrogenase complex; NOX4: Nicotinamide adenine dinucleotide phosphate hydrogen oxidase 4; ROS: Reactive oxygen species; FAP: fibroblast activation protein.

**Figure S6:**
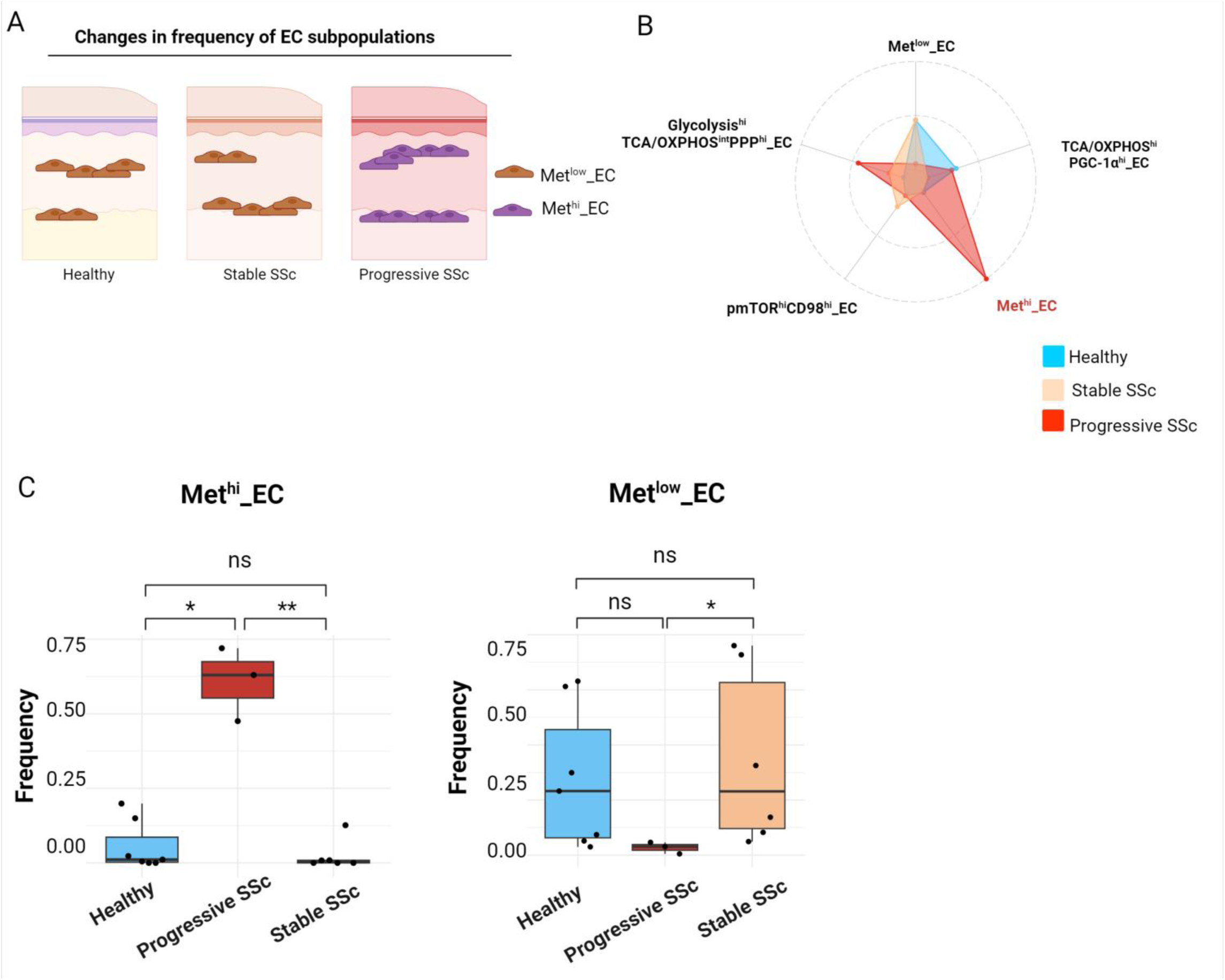
Shifts in frequencies of metabolically defined endothelial cell subpopulations in SSc patients with progressive skin fibrosis. **A.** Illustration of the shifts in endothelial cell subsets across the groups: progressive SSc, stable SSc or controls. **B.** Radar chart illustrating the shifts in frequencies of different End-MET subpopulations (normalized for each population) across the groups: progressive SSc, stable SSc or controls. The Met^hi^_EC are highlighted in red as significantly increased in progressive SSc. **C-D.** Frequencies of Met^hi^_EC or Met^low^_EC across the groups: progressive SSc, stable SSc or controls, shown as box plots. Statistical significance was determined by the Kruskal Wallis test with Dunn’s post-hoc test. 0.05>*P*>0.01*, 0.01>*P*>0.001**, *P*<0.001***. This figure was created with Biorender.com. UMAP: Uniform Manifold Approximation and Projection; EC: endothelial cells; End-MET: metabolically defined endothelial cell subsets; SSc: systemic sclerosis.

**Figure S7:**
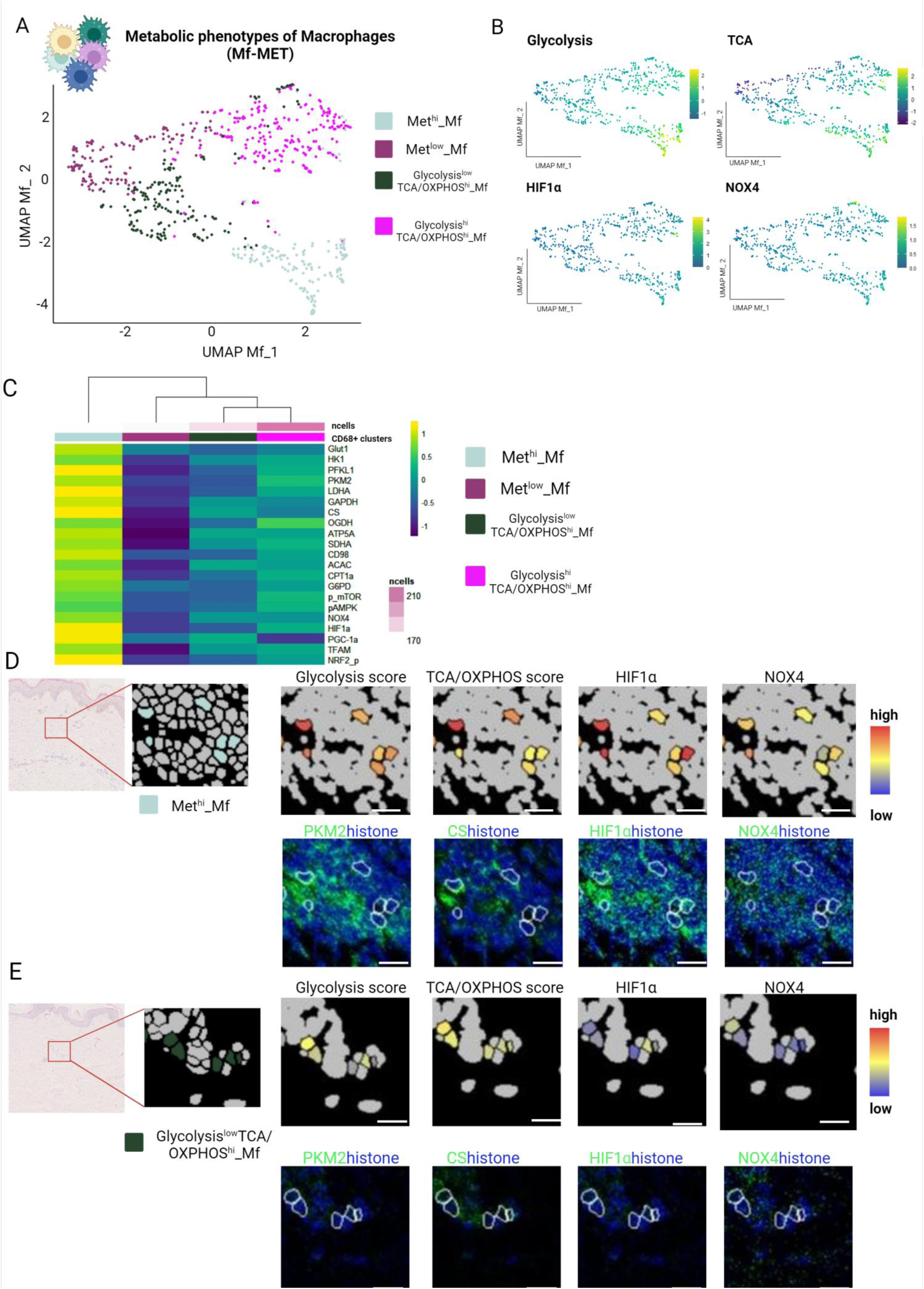
Identification of metabolically defined populations of macrophages. **A.** UMAP plots generated based on expression of metabolic markers and colour coded by metabolically defined macrophage subsets, and illustration of the distinct macrophage subsets. **B.** Glycolysis score (calculated by averaging the expression of enzymes belonging to the glycolysis pathway: GLUT1, HK1, PFKL1, PKM2, LDHA, GAPDH), TCA/OXPHOS score (calculated by averaging the expression of enzymes belonging to the TCA cycle and OXPHOS: CS, OGDH, ATP5A, SDHA), hypoxia (expression of HIF1α) and ROS (expression of NOX4) signaling in macrophages plotted on UMAP. **C.** Expression of metabolic markers across metabolically defined macrophage subsets, plotted as heatmap. **D-E.** Representative images illustrating spatial distribution of the Met^hi^_Mf (D) and of Glycolysis^low^TCA/OXPHOS^hi^_Mf (E), along glycolysis and TCA/ETC scores and hypoxia and ROS signaling, plotted on cell segmentation masks, and images showing expression of PKM2 (a glycolytic enzyme), CS (an enzyme from the TCA cycle), HIF1α (hypoxia signaling) and NOX4 (ROS signaling) at pixel level. Corresponding HE images were included. Scale bars: 25 µm. This figure was created with Biorender.com. UMAP: Uniform Manifold Approximation and Projection; HE; hematoxylin and eosin; Mf: macrophages; TCA: tricarboxylic acid; OXPHOS: oxidative phosphorylation; GLUT1: Glucose transporter 1; HK1: Hexokinase1; PFKL1: Phosphofructokinase Ligand 1; PKM2: Pyruvate kinase isoform M2; LDHA: Lactate dehydrogenase A; GAPDH: Glyceraldehyde 3-phosphate dehydrogenase; CS: Citrate synthase; OGDH: 2-oxoglutarate dehydrogenase; ATP5A: Mitochondrial adenosine triphosphatase alpha chain; SDHA: succinate dehydrogenase complex; NOX4: Nicotinamide adenine dinucleotide phosphate hydrogen oxidase 4; ROS: Reactive oxygen species.

**Figure S8:**
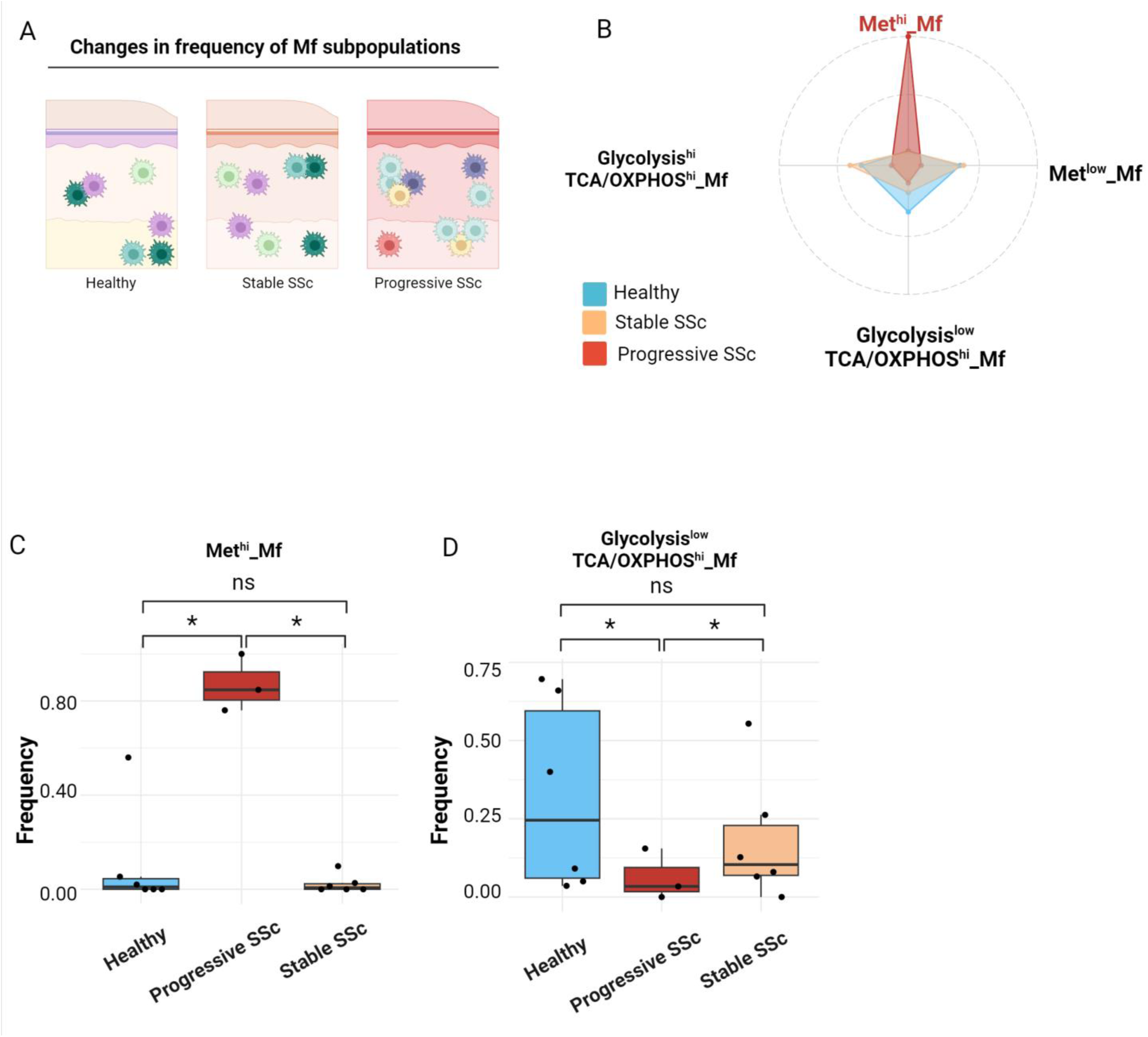
Shifts in frequencies of metabolically defined macrophage subpopulations in SSc patients with progressive skin fibrosis. **A.** Illustration of the shifts in fibroblast subsets across the groups: progressive SSc, stable SSc or controls. **B.** Radar chart illustrating the shifts in frequencies of different Mf-MET subpopulations (normalized for each population) across the groups: progressive SSc, stable SSc or controls. The Met^hi^_Mf are highlighted in red as significantly increased in progressive SSc. **C-D.** Frequencies of Met^hi^_Mf or Glycolysis^low^TCA/OXPHOS^hi^_Mf across the groups: progressive SSc, stable SSc or controls, shown as box plots. Statistical significance was determined by the Kruskal Wallis test with Dunn’s post-hoc test. 0.05>*P*>0.01*, 0.01>*P*>0.001**, *P*<0.001***. This figure was created with Biorender.com. UMAP: Uniform Manifold Approximation and Projection; Mf: macrophages; Mf-MET: metabolically defined macrophage subsets; SSc: systemic sclerosis.

**Figure S9:**
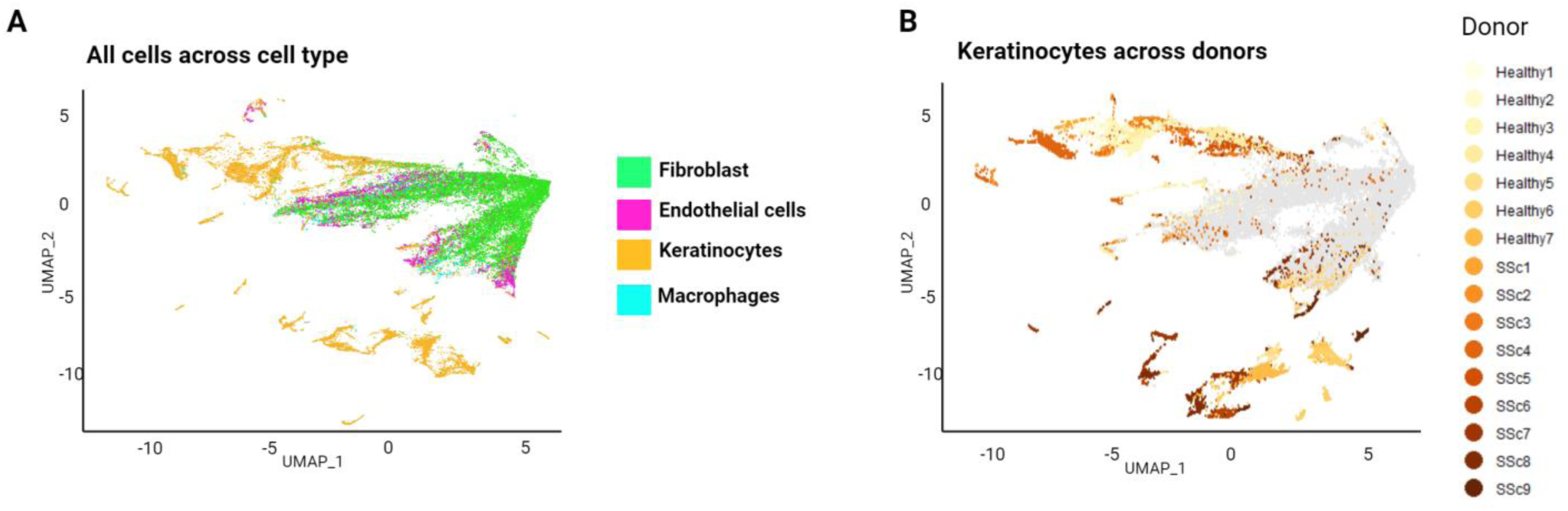
Donor-to-donor variation of metabolic regulome in keratinocytes. **A.** UMAP plots generated based on expression of metabolic markers and colour coded by fibroblasts, keratinocytes, endothelial cells and macrophages. **B.** UMAP plots generated based on expression of metabolic markers and with keratinocytes colour coded by donor, and other cells colored in gray. UMAP: Uniform Manifold Approximation and Projection.

**Figure S10:**
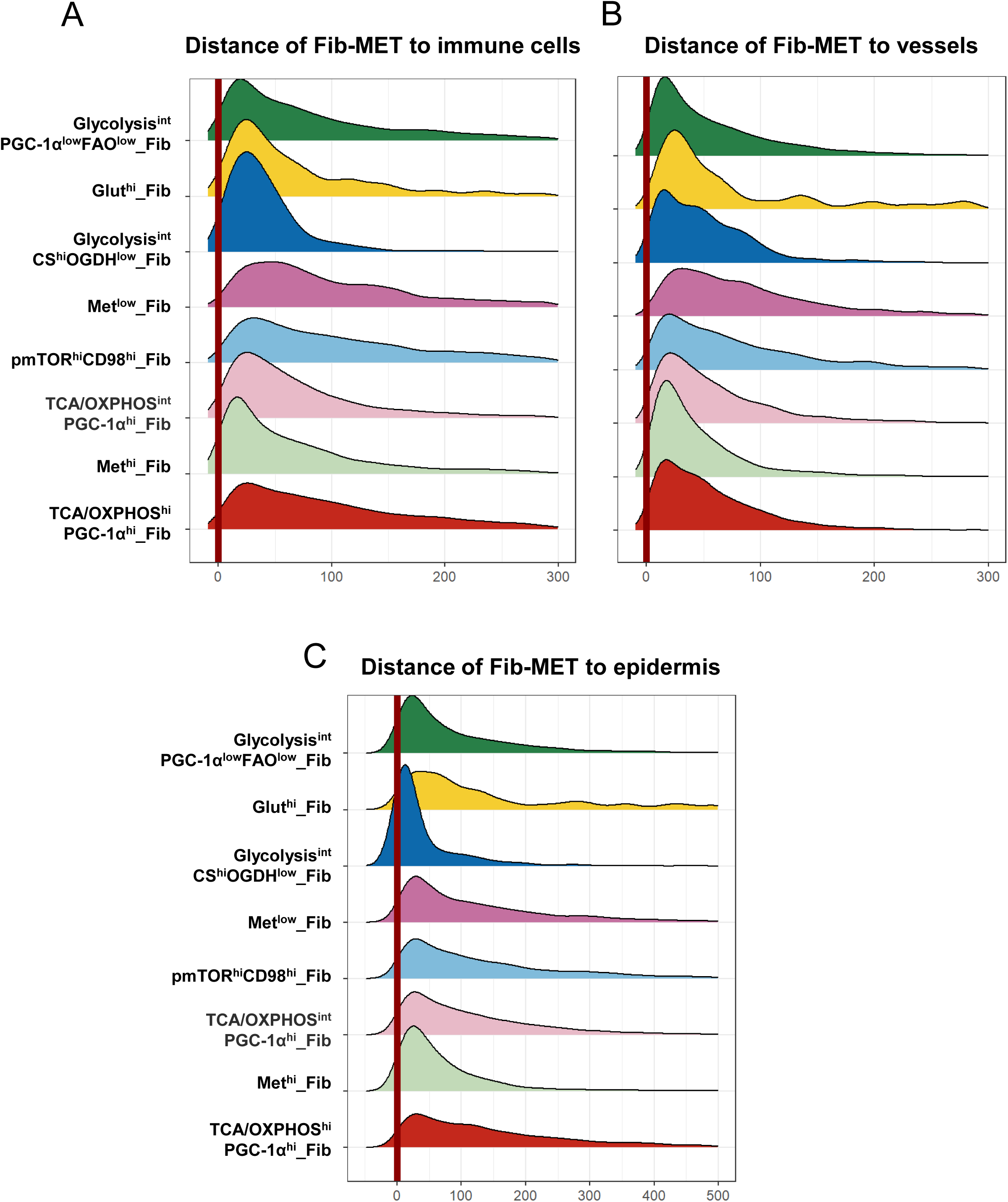
Distances of distinct metabolically defined fibroblast subsets to epidermis, vessels or immune cells A-C. Ridge plots showing the distance of Fib-MET subsets to immune cells (A), to vessels (B) or to epidermis (C). Fib-MET: metabolically defined fibroblast subsets.

**Figure S11:**
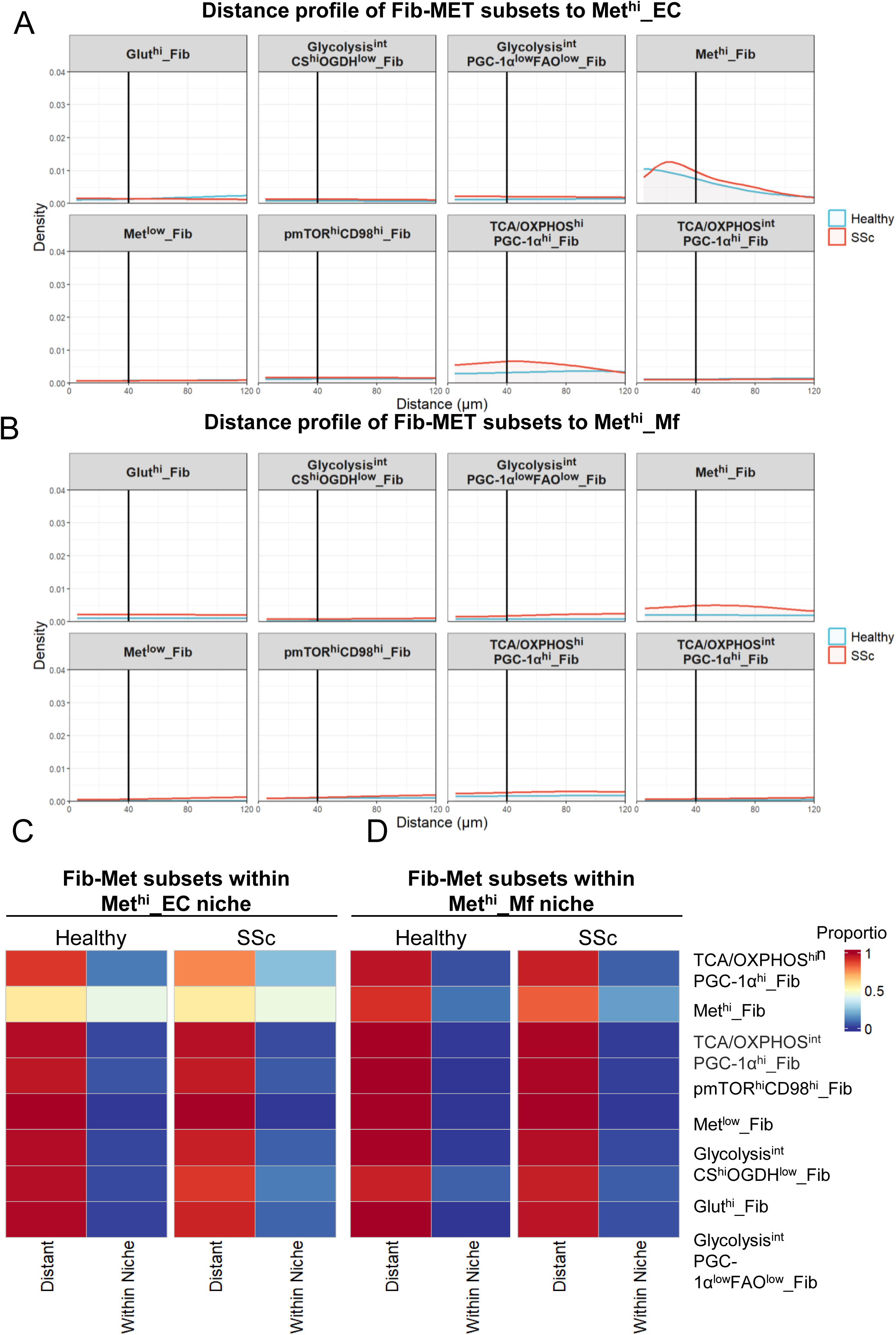
**Spatial relationships of metabolically defined fibroblast subsets with Met^hi^_EC or Met^hi^_Mf A-B**. Distance of Fib-MET subsets to Met^hi^_EC (A) or Met^hi^_Mf (B) in SSc patients and controls shown as density plots. **C-D.** Proportions of Fib-MET subsets present within the Met^hi^_EC niche (C) or Met^hi^_Mf niche (D) or outside the respective niches in SSc patients or controls, shown as heatmaps. SSc: systemic sclerosis; EC: endothelial cells; Mf: macrophages Fib-MET: metabolically defined fibroblast subsets.

**Figure S12:**
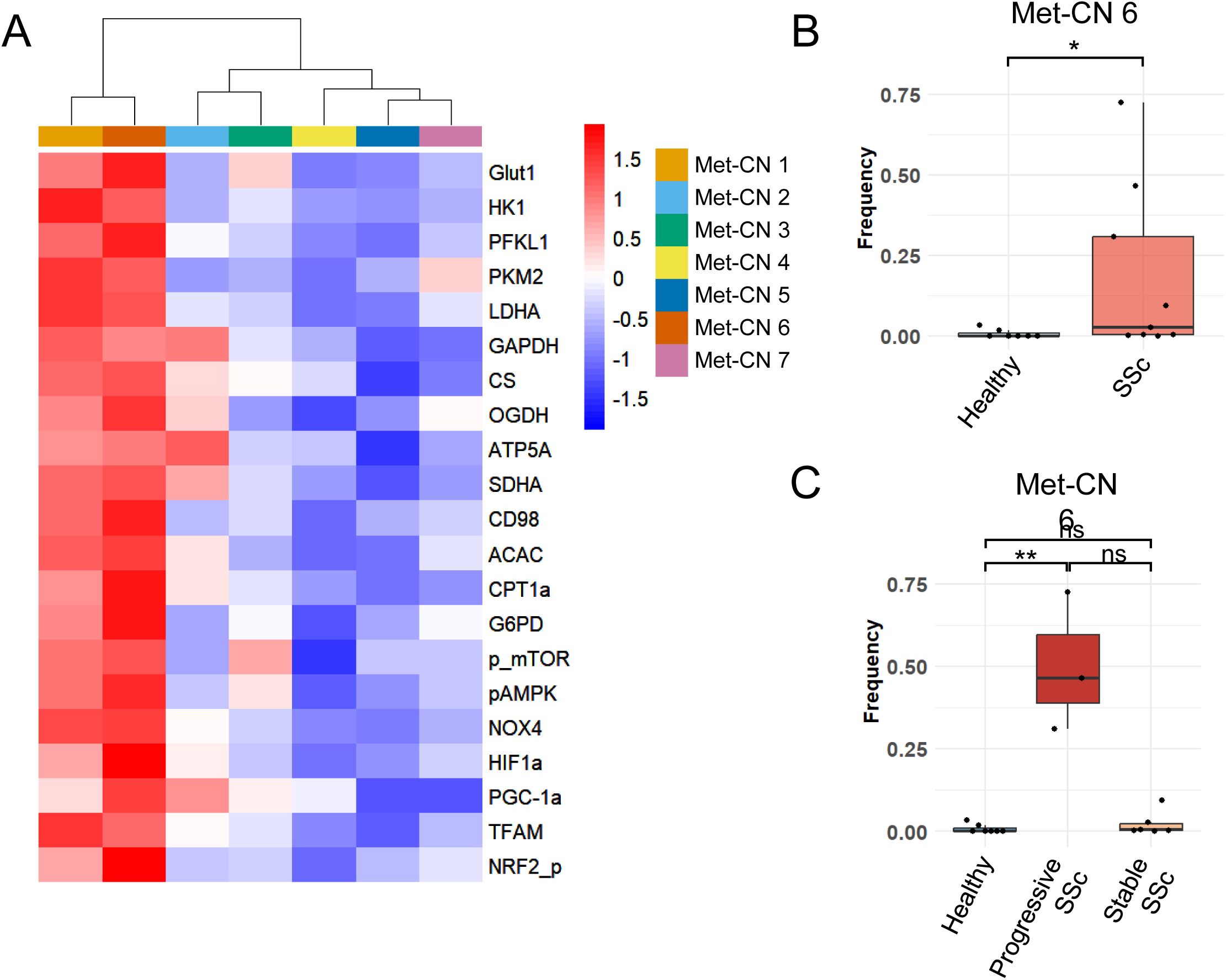
Expression of metabolic markers across metabolically defined cellular neighbourhoods and changes in the frequencies of these neighbourhoods in progressive SSc. **A.** Expression of metabolic markers across different metabolically defined cellular neighbourhoods, shown as heatmap. **B-C.** Frequency of Met-CN6 (among the Met-CNs) across the conditions: SSc or controls (B), or progressive SSc, stable SSc or controls (C). SSc: systemic sclerosis; Met-CN: metabolically defined cellular neighbourhood.

**Figure S13:**
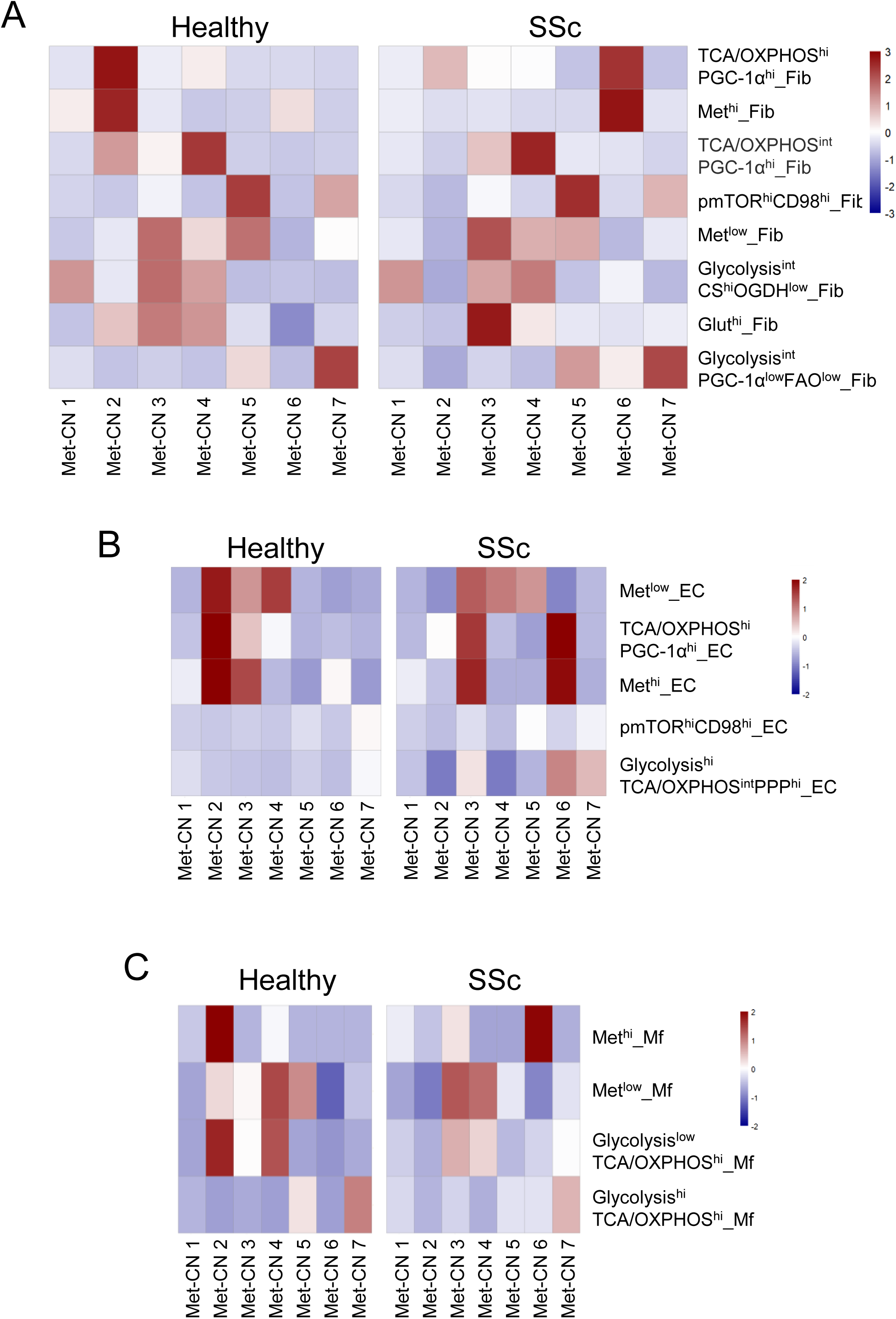
Shifts in composition of metabolically defined cellular neighbourhoods in SSc patients A-C. Heatmaps showing the shifts in Fib-MET (A), End-MET (B) or Mf-MET (C) subsets composition of metabolically defined cellular neighbourhoods in SSc compared to controls. SSc: systemic sclerosis; Fib-MET: metabolically defined fibroblast subsets; End-MET: metabolically defined endothelial cell subsets; Mf-MET: metabolically defined macrophage subsets.

**Figure S14:**
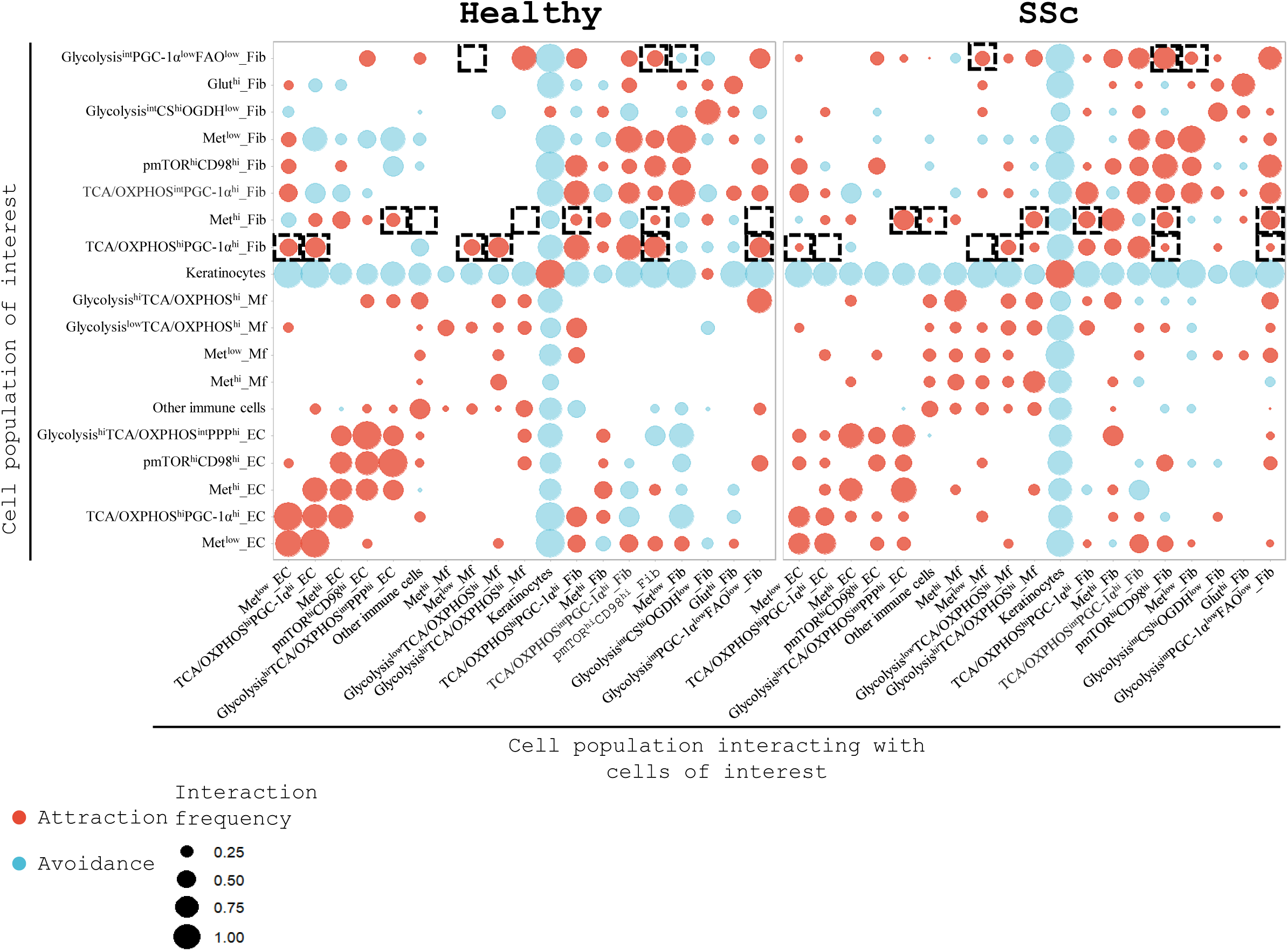
Cell-cell interactions between metabolically defined fibroblast, endothelial cell and macropahge subsets in SSc patients and controls. Dot plot highlighting the changes in interaction frequencies, i.e. proportion of samples with statistically significant interactions, between Fib-MET, End-MET or Mf-MET subsets in SSc patients compared to controls. The size of the dot represents the interaction frequency. Cellular attraction (red) and avoidance (blue) are indicated in the plots. SSc: systemic sclerosis; Fib-MET: metabolically defined fibroblast subsets; End-MET: metabolically defined endothelial cell subsets; Mf-MET: metabolically defined macrophage subsets.

**Figure S15:**
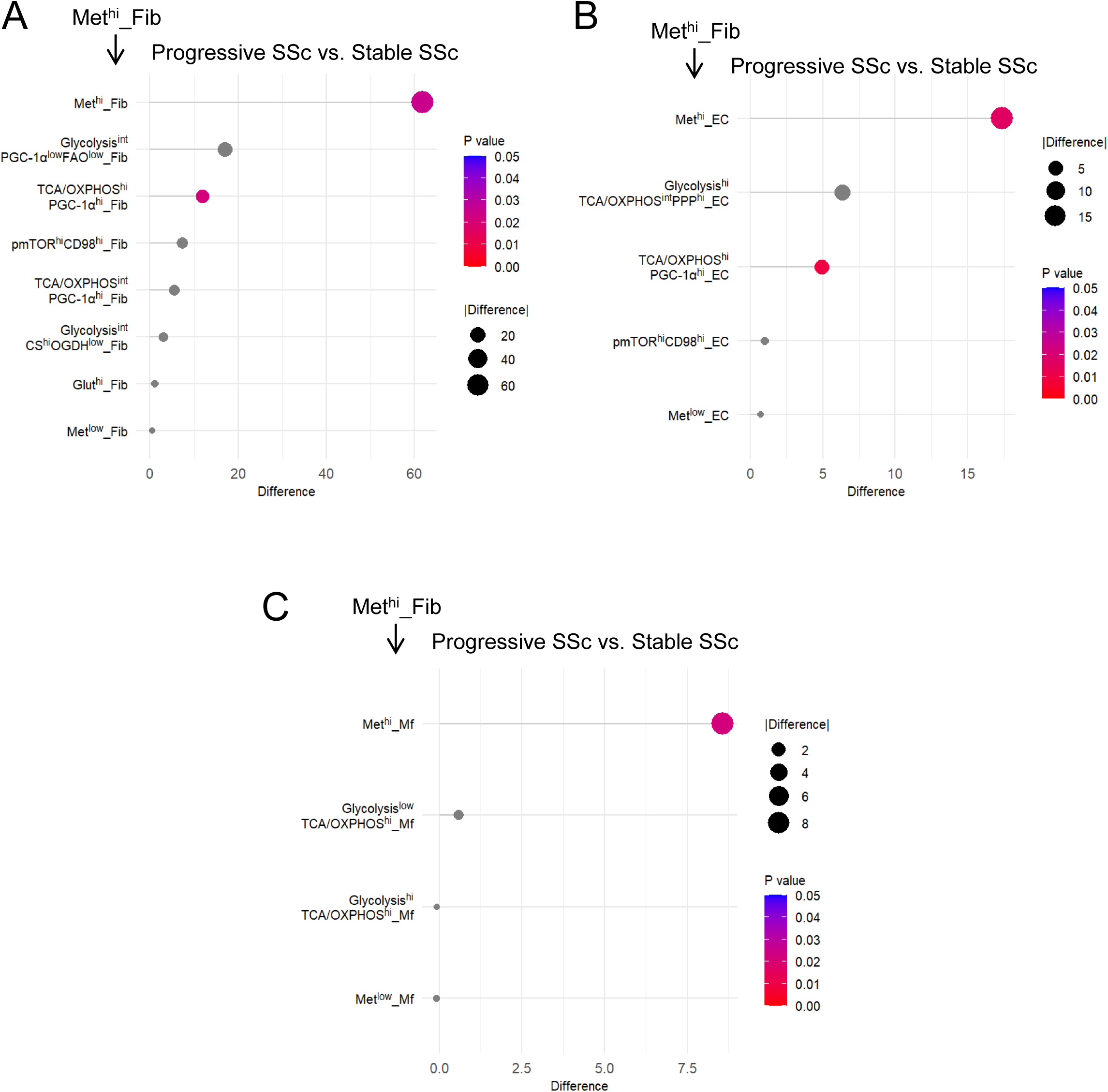
Changes in frequencies of Met^hi^_Fib in spatial proximity to individual Fib-MET, End-MET or Mf-MET subsets in progressive SSc A-C. Dot plot illustrating the difference in frequencies of Met^hi^_Fib in spatial proximity to individual Fib-MET subsets (A), End-MET subsets (B) or Mf-MET subsets (C) (among all fibroblasts) in progressive vs. stable SSc. The size of the dots represents the module of the difference in frequency, and the color represents the p-value. Statistical significance was determined by the Wilcoxon test. SSc: systemic sclerosis; Fib-MET: metabolically defined fibroblast subsets; End-MET: metabolically defined endothelial subsets; Mf-MET: metabolically defined macrophage subsets; Fib: fibroblasts.

**Figure S16:**
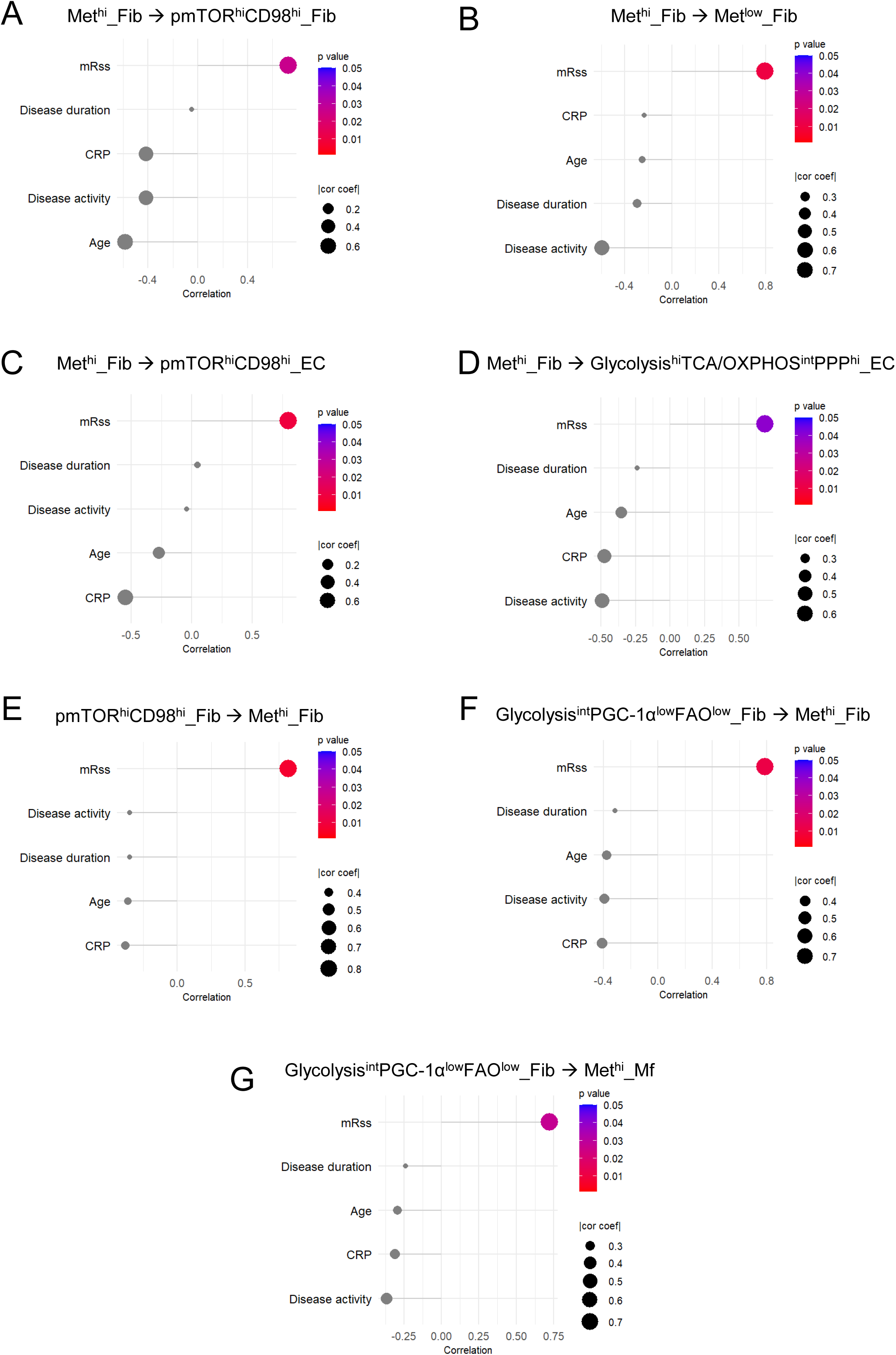
Other clinical parameters except mRSS do not correlate with the frequencies of metabolically defined fibroblast subsets in spatial proximity to distinct End-MET or Mf-MET subsets in SSc A-G. Dot plot illustrating the correlation of mRSS, disease duration, CRP, disease activity and age with the frequency of Met^hi^_Fib in spatial proximity to pmTOR^hi^CD98^hi^_Fib (A), Met^low^_Fib (B), pmTOR^hi^CD98^hi^_EC (C), Glycolysis^hi^TCA/OXPHOS^int^PPP^hi^_EC (D), or of pmTOR^hi^CD98^hi^_Fib in spatial proximity to Met^hi^_Fib (E), or of Glycolysis^int^PGC-1α^low^FAO^low^_Fib in spatial proximity to Met^hi^_Fib (F) or Met^hi^_Mf (G) (among all fibroblasts). The size of the dots represents the coefficient of correlation, and the color represents the p-value. Statistical significance was determined by Spearman’s correlation analysis. SSc: systemic sclerosis; End-MET: metabolically defined endothelial subsets; Mf-MET: metabolically defined macrophage subsets; Fib: fibroblasts, EC: endothelial cells; Mf: macrophages; mRSS: modified Rodnan skin score; TCA: tricarboxylic acid; OXPHOS: oxidative phosphorylation; FAO: fatty acid oxidation.

**Figure S17:**
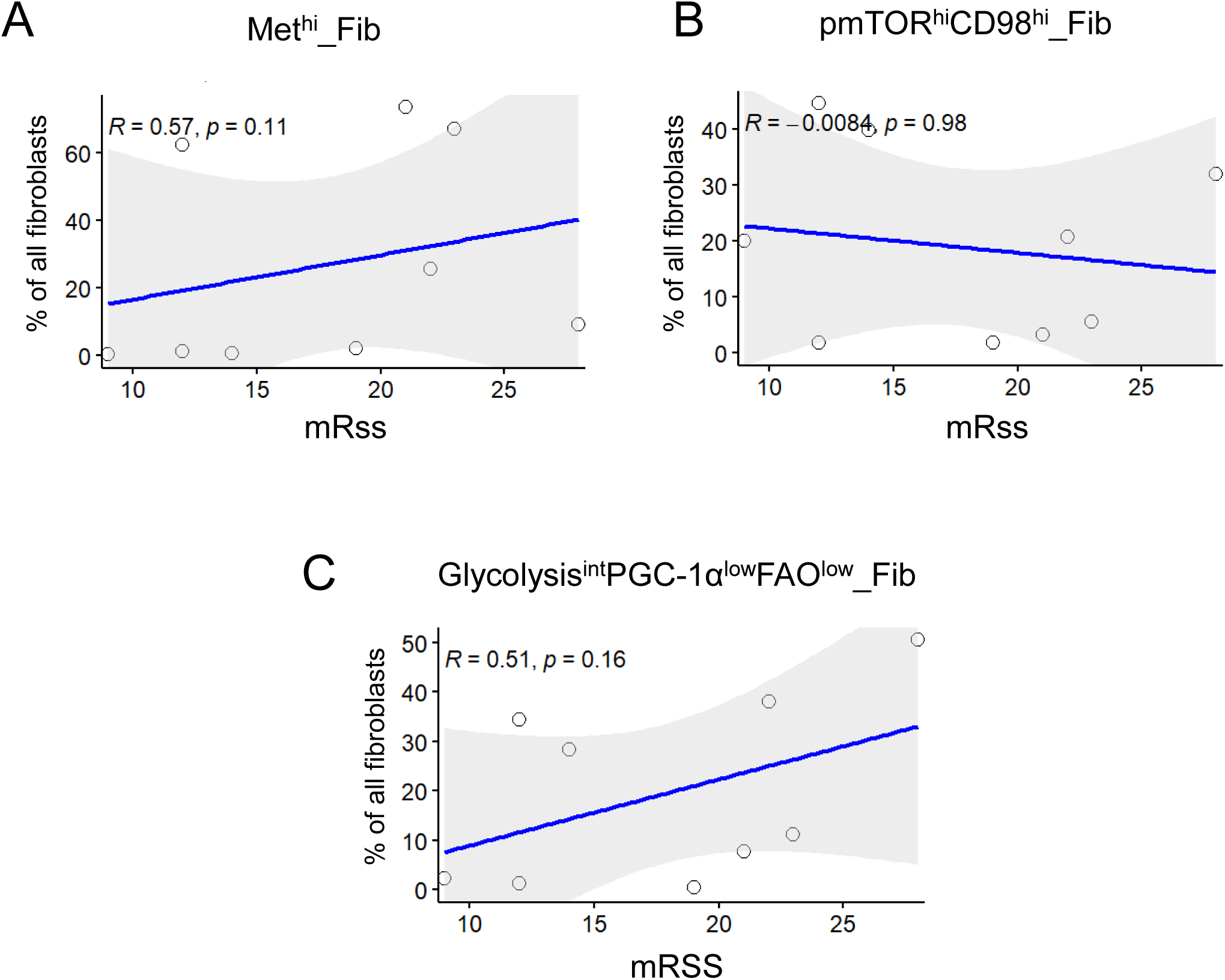
The frequencies of metabolically defined fibroblast subsets do not correlate with mRSS when not accounting for their interactions A-C. Scatter plots illustrating the correlation between mRSS and the frequency of all Met^hi^_Fib (A), of pmTOR^hi^CD98^hi^_Fib (B), or of Glycolysis^int^PGC-1α^low^FAO^low^_Fib (C), regardless of their interactions (among all fibroblasts). Statistical significance was determined by Spearman’s correlation analysis. Linear regression lines with the 95% CI, the Spearman’s rank correlation coefficient and the p-value are included in the scatter plots. Fib: fibroblasts; mRSS: modified Rodnan skin score; TCA: tricarboxylic acid; OXPHOS: oxidative phosphorylation; FAO: fatty acid oxidation.

**Figure S18:**
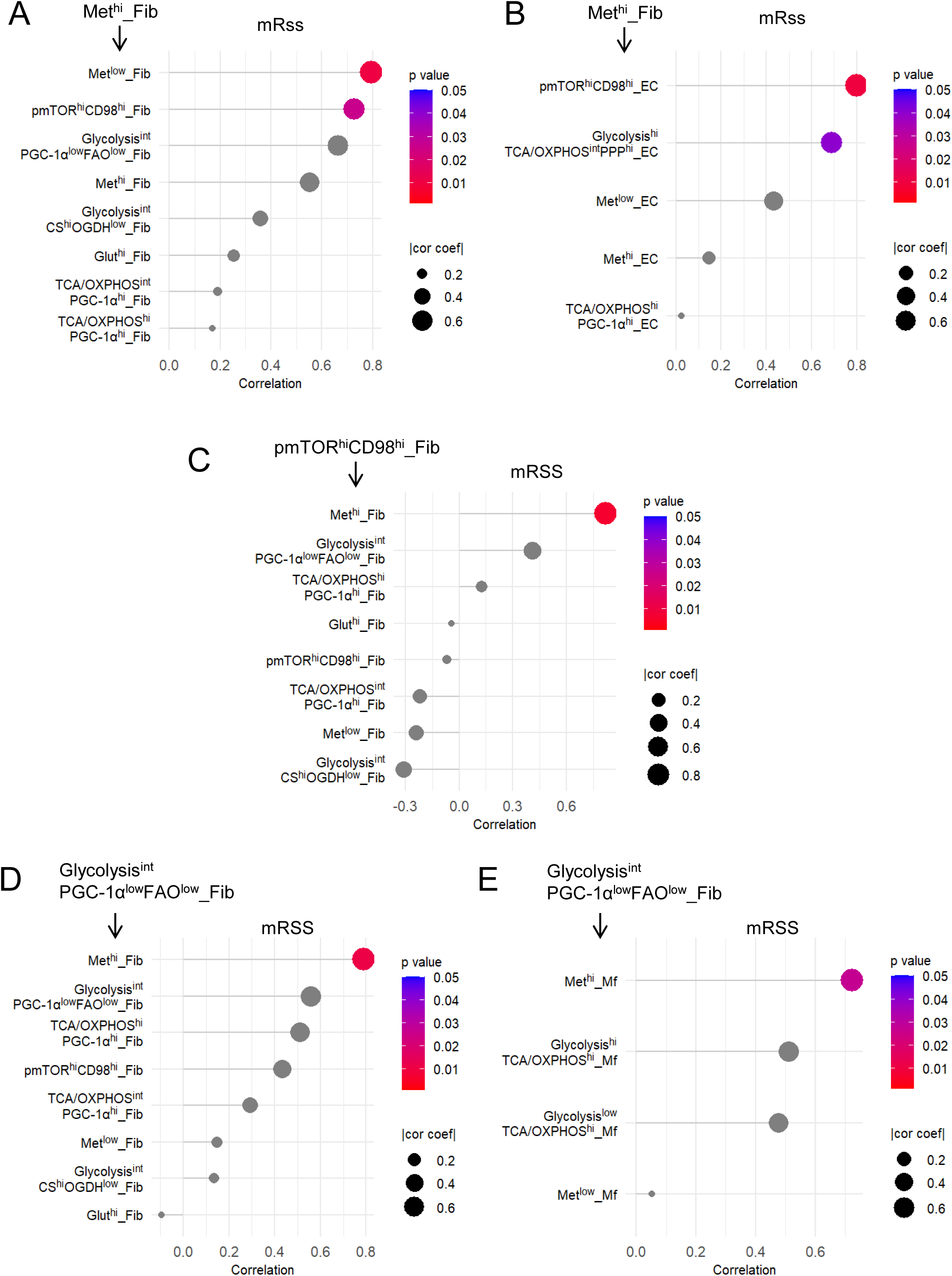
Correlations of mRSS with specific interactions between metabolically defined cell subsets A-E. Dot plot illustrating the correlation of mRSS with the frequency of Met^hi^_Fib in spatial proximity to individual Fib-MET subsets (A) or End-MET subsets (B), or of pmTOR^hi^CD98^hi^_Fib in spatial proximity to individual Fib-MET subsets (C), or of Glycolysis^int^PGC-1α^low^FAO^low^_Fib in spatial proximity to individual Fib-MET subsets (D) or Mf-MET subsets (E) (among all fibroblasts). The size of the dots represents the coefficient of correlation, and the color represents the p-value. Statistical significance was determined by Spearman’s correlation analysis. Fib-MET: metabolically defined fibroblast subsets; End-MET: metabolically defined endothelial subsets; Mf-MET: metabolically defined macrophage subsets; Fib: fibroblasts; mRSS: modified Rodnan skin score; TCA: tricarboxylic acid; OXPHOS: oxidative phosphorylation; FAO: fatty acid oxidation.

**Table S1.**

| **Metal** | **Antigen** | **Clone** | **Company** | **Catalog number** | **Dilution** |
| --- | --- | --- | --- | --- | --- |
| 113In | Glut1 | EPR3915 | Abcam | ab252403 | 1/200 |
| 141Pr | PKM2 | D78A4 | Cell Signaling | 95251 | 1/200 |
| 142Nd | CD98 | D3F9D | Cell Signaling | 81977SF | 1/100 |
| 143Nd | P_mTOR | EPR426(2) | Abcam | ab177734 | 1/200 |
| 144Nd | P_NRF2 | EP1809Y | Abcam | ab18084 | 1/50 |
| 145Nd | CD31 | Polyclonal | R&D | AF3628 | 1/200 |
| 146Nd | G6PD | EPR20668 | Abcam | ab23182 | 1/200 |
| 147Sm | mtTFA | C-9 | Santa Cruz | sc-376672 L | 1/200 |
| 148Nd | PDGFRα | Polyclonal | R&D | AF307NA | 1/200 |
| 149Sm | LDHA | EP1566Y | Abcam | ab21959 | 1/50 |
| 150Nd | α-SMA | ASM-1/1A4 | Sigma Aldrich | CBL171-I | 1/200 |
| 151Eu | SDHA | 2E3GC12FB<br>2AE2 | Abcam | ab14715 | 1/200 |
| 152Sm | CD45 | D9M8I | SBT/<br>Fluidigm | 3152018D | 1/200 |
| 153Eu | CD20 | H1 | BD | 555677 | 1/800 |
| 154Sm | p_ribosom.S6 | Ser235/236 | Cell signaling | 81736 | 1/400 |
| 155Gd | p_AMPK | (Thr172)<br>(40H9) | Cell signaling | 74281 | 1/200 |
| 156Gd | PGC-1α | Polyclonal | Invitrogen | PA5-38022 | 1/200 |
| 158Gd | E-Cadherin | 24E10 | Fluidigm | 3158029 | 1/200 |
| 159Tb | OGDH | Polyclonal | Invitrogen | PA5-28195 | 1/200 |
| 160Gd | Podoplanin | LpMab-21 | Biolegend | 395002 | 1/200 |
| 161Dy | GAPDH | 6C5 | Abcam | ab8245 | 1/200 |
| 162Dy | Cdh11 | 23C6 | - | Provided by<br>M.B. Brenner | 1/200 |
| 163Dy | COL1 | EPR7785 | Abcam | ab215969 | 1/100 |
| 164Dy | CS | EPR8067 | Abcam | ab233838 | 1/100 |
| 165Ho | HK1 | EPR10134(B<br>) | Abcam | ab233837 | 1/800 |
| 166Er | ATP5A | 15H4C4 | Abcam | ab14748 | 1/200 |
| 167Er | PFKL1+PFK<br>M | EPR11904 | Abcam | ab240237 | 1/200 |
| 168Er | Ki-67 | (B56) | Fluidigm | 3168022D | 1/400 |
| 169Tm | CPT1a | 8F6AE9 | Abcam | ab128568 | 1/200 |
| 170Er | CD3 | Polyclonal | SBT/<br>Fluidigm | 3170019D | 1/200 |
| 171Yb | ACAC | EPR4971 | Abcam | ab231681 | 1/200 |
| 172Yb | NOX4 | Polyclonal | Abcam | ab305657 | 1/50 |
| 173Yb | HIF1a | Polyclonal | R&D | AF1935 | 1/50 |
| 174Yb | FAP | Polyclonal | R&D | AF3715 | 1/200 |
| 175Lu | CD68 | KP1 | Biolegend | 916104 | 1/800 |
| 176Yb | Histone 3 | D1H2 | SBT/<br>Fluidigm | 4276023 | 1/1600 |
| 191Ir/193Ir | Iridium | NA | SBT/<br>Fluidigm | 201192A | 1/400 |
| 195Pt | Segmentation<br>kit (ICSK1) | NA | SBT/<br>Fluidigm | Tis-00001 | 1/400 |
| 196Pt | Segmentation<br>kit (ICSK2) | NA | SBT/<br>Fluidigm | Tis-00001 | 1/400 |

**Table S2.** Demographic and clinical characteristics of the SSc patients.

| **Characteristics** | **SSc patients (number (percentage) or median [IQR])** |
| --- | --- |
| Gender (male / female) | 5 / 4 (55.5% / 44.4%) |
| Age (years) | 54.0 [50.0 – 66.0] |
| SSc subtype (lcSSc / dcSSc) | 2 / 7 (22.2% / 77.7%) |
| Disease duration from first non-RP (months) | 84.0 [24.0 – 132.0] |
| Patients with early SSc | 7 (77.7%) |
| Modified Rodnan skin score (mRSS) | 19.0 [12.0 – 22.0] |
| Patients with progressive skin fibrosis | 3 (33.3%) |
| EUSTAR SSc activity score | 5.25 [4 – 6.25] |
| Patients with history of digital ulcer(s) (DU) | 7 (77.7%) |
| Arthralgia | 3 (33.3%) |
| Interstitial lung disease (ILD) | 4 (44.4%) |
| Pulmonary hypertension (PH) | 0 (0%) |
| Tendon friction rub(s) (TFR) | 1 (11.1%) |
| CRP (mg/L) | 6.4 [4.9 – 11.4] |
| Positive for anti Scl-70 autoantibody | 3 (33.3%) |
| Positive for anti-RNA polymerase III autoantibody | 4 (44.4%) |
| Positive for anti-centromere B autoantibody | 2 (22.2%) |

## References

1. Gabrielli, A., Avvedimento, E.V. & Krieg, T. Scleroderma. N Engl J Med 360, 1989–2003 (2009).

2. Distler, J.H.W., et al. Shared and distinct mechanisms of fibrosis. Nat Rev Rheumatol 15, 705–730 (2019).

3. Gyorfi, A.H., Matei, A.E. & Distler, J.H.W. Targeting TGF-beta signaling for the treatment of fibrosis. Matrix Biol 68-69, 8-27 (2018).

4. Garrett, S.M., Baker Frost, D. & Feghali-Bostwick, C. The mighty fibroblast and its utility in scleroderma research. J Scleroderma Relat Disord 2, 69–134 (2017).

5. Hinz, B. & Lagares, D. Evasion of apoptosis by myofibroblasts: a hallmark of fibrotic diseases. Nat Rev Rheumatol 16, 11–31 (2020).

6. Manetti, M., et al. Endothelial-to-mesenchymal transition contributes to endothelial dysfunction and dermal fibrosis in systemic sclerosis. Ann Rheum Dis 76, 924–934 (2017).

7. Jimenez, S.A. & Piera-Velazquez, S. Endothelial to mesenchymal transition (EndoMT) in the pathogenesis of Systemic Sclerosis-associated pulmonary fibrosis and pulmonary arterial hypertension. Myth or reality? Matrix Biol 51, 26–36 (2016).

8. Di Benedetto, P., et al. Endothelial-to-mesenchymal transition in systemic sclerosis. Clin Exp Immunol 205, 12–27 (2021).

9. Rius Rigau, A., et al. Characterization of Vascular Niche in Systemic Sclerosis by Spatial Proteomics. Circ Res 134, 875–891 (2024).

10. Al-Adwi, Y., et al. Macrophages as determinants and regulators of fibrosis in systemic sclerosis. Rheumatology (Oxford) 62, 535–545 (2023).

11. Bhandari, R., et al. Profibrotic Activation of Human Macrophages in Systemic Sclerosis. Arthritis Rheumatol 72, 1160–1169 (2020).

12. Feng, L., et al. Immunometabolism changes in fibrosis: from mechanisms to therapeutic strategies. Front Pharmacol 14, 1243675 (2023).

13. Ung, C.Y., Onoufriadis, A., Parsons, M., McGrath, J.A. & Shaw, T.J. Metabolic perturbations in fibrosis disease. Int J Biochem Cell Biol 139, 106073 (2021).

14. Henderson, J. & O’Reilly, S. The emerging role of metabolism in fibrosis. Trends Endocrinol Metab 32, 639–653 (2021).

15. Zeng, H., et al. Suppression of PFKFB3-driven glycolysis restrains endothelial-to-mesenchymal transition and fibrotic response. Signal Transduct Target Ther 7, 303 (2022).

16. Andreucci, E., et al. Glycolysis-derived acidic microenvironment as a driver of endothelial dysfunction in systemic sclerosis. Rheumatology (Oxford) 60, 4508–4519 (2021).

17. Cantanhede, I.G., et al. Exploring metabolism in scleroderma reveals opportunities for pharmacological intervention for therapy in fibrosis. Front Immunol 13, 1004949 (2022).

18. Hartmann, F.J., et al. Single-cell metabolic profiling of human cytotoxic T cells. Nat Biotechnol 39, 186–197 (2021).

19. Gyorfi, A.H., et al. Engrailed 1 coordinates cytoskeletal reorganization to induce myofibroblast differentiation. J Exp Med 218(2021).

20. Rius Rigau, A., et al. Imaging mass cytometry-based characterisation of fibroblast subsets and their cellular niches in systemic sclerosis. Ann Rheum Dis (2024).

21. Windhager, J., et al. An end-to-end workflow for multiplexed image processing and analysis. Nat Protoc 18, 3565–3613 (2023).

22. Greenwald, N.F., et al. Whole-cell segmentation of tissue images with human-level performance using large-scale data annotation and deep learning. Nat Biotechnol 40, 555–565 (2022).

23. Righelli, D., et al. SpatialExperiment: infrastructure for spatially-resolved transcriptomics data in R using Bioconductor. Bioinformatics 38, 3128–3131 (2022).

24. Eling, N., Damond, N., Hoch, T. & Bodenmiller, B. cytomapper: an R/Bioconductor package for visualization of highly multiplexed imaging data. Bioinformatics 36, 5706–5708 (2021).

25. Rigau AR, L.Y.-N., Matei A-E, Györfi A-H, Bruch P-M, Koziel S, Gabrielli A, Kreuter A, Wang J, Dietrich S, Schett G, Distler JHW, Liang M.. Characterization of vascular niche in systemic sclerosis by spatial proteomics. Circ Res. (2024).

26. Li, Y.-N., et al. Spatially informed phenotyping by cyclic-in-situ-hybridization identifies novel fibroblast populations and their pathogenic niches in systemic sclerosis. *bioRxiv*, 2024.2012.2028.630505 (2024).

27. Piera-Velazquez, S. & Jimenez, S.A. Endothelial to Mesenchymal Transition: Role in Physiology and in the Pathogenesis of Human Diseases. Physiol Rev 99, 1281–1324 (2019).

28. He, X., et al. Intimate intertwining of the pathogenesis of hypoxia and systemic sclerosis: A transcriptome integration analysis. Front Immunol 13, 929289 (2022).

29. Svegliati, S., Spadoni, T., Moroncini, G. & Gabrielli, A. NADPH oxidase, oxidative stress and fibrosis in systemic sclerosis. Free Radic Biol Med 125, 90–97 (2018).

30. Tabib, T., et al. Myofibroblast transcriptome indicates SFRP2(hi) fibroblast progenitors in systemic sclerosis skin. Nat Commun 12, 4384 (2021).

31. Ma, F., et al. Systems-based identification of the Hippo pathway for promoting fibrotic mesenchymal differentiation in systemic sclerosis. Nat Commun 15, 210 (2024).

32. Gur, C., et al. LGR5 expressing skin fibroblasts define a major cellular hub perturbed in scleroderma. Cell 185, 1373–1388 e1320 (2022).

33. Gibb, A.A., Lazaropoulos, M.P. & Elrod, J.W. Myofibroblasts and Fibrosis: Mitochondrial and Metabolic Control of Cellular Differentiation. Circ Res 127, 427–447 (2020).

34. Xie, N., et al. Glycolytic Reprogramming in Myofibroblast Differentiation and Lung Fibrosis. Am J Respir Crit Care Med 192, 1462–1474 (2015).

35. Ung, C.Y., Onoufriadis, A., Parsons, M., McGrath, J.A. & Shaw, T.J. Metabolic perturbations in fibrosis disease. The International Journal of Biochemistry & Cell Biology 139, 106073 (2021).

36. Li, X., et al. Mitochondrial dysfunction in fibrotic diseases. Cell Death Discovery 6, 80 (2020).

37. Zhou, X., et al. Impaired Mitochondrial Transcription Factor A Expression Promotes Mitochondrial Damage to Drive Fibroblast Activation and Fibrosis in Systemic Sclerosis. Arthritis & Rheumatology 74, 871–881 (2022).

38. Soldano, S., et al. Increase in circulating cells coexpressing M1 and M2 macrophage surface markers in patients with systemic sclerosis. Annals of the Rheumatic Diseases 77, 1842 (2018).

39. Andreucci, E., et al. Glycolysis-derived acidic microenvironment as a driver of endothelial dysfunction in systemic sclerosis. Rheumatology 60, 4508–4519 (2021).

